# Polymodal Allosteric Regulation of Type 1 Serine/Threonine Kinase Receptors via a Conserved Electrostatic Lock

**DOI:** 10.1101/170837

**Authors:** Wesley M. Botello-Smith, Abdelaziz Alsamarah, Payal Chatterjee, Chen Xie, Jerome J. Lacroix, Jijun Hao, Yun Luo

## Abstract

Type 1 Serine/Threonine Kinase Receptors (STKR1) transduce a wide spectrum of biological signals mediated by TGF-β superfamily members. The STKR1 activity is tightly controlled by their regulatory glycine-serine rich (GS) domain adjacent to the kinase domain. Despite decades of studies, it remains unknown how physiological or pathological GS domain modifications are coupled to STKR1 kinase activity. Here, by performing molecular dynamics simulations and free energy calculation of Activin-Like Kinase 2 (ALK2), we found that GS domain phosphorylation, FKBP12 dissociation, and disease mutations all destabilize a D354-R375 salt-bridge, which normally acts as an electrostatic lock to prevent coordination of adenosine triphosphate (ATP) to the catalytic site. We developed a WAFEX-guided principal analysis and unraveled how phosphorylation destabilizes this highly conserved salt-bridge in temporal and physical space. Using current-flow betweenness scores, we identified an allosteric network of residue-residue contacts between the GS domain and the catalytic site that controls the formation and disruption of this salt bridge. Importantly, our novel network analysis approach revealed how certain disease-causing mutations bypass FKBP12-mediated kinase inhibition to produce leaky signaling in the absence of ligand. We further provide experimental evidence that this salt-bridge lock exists in other STKR1s, and acts as a general safety mechanism in STKR1 to prevent pathological leaky signaling. In summary, our study provides a compelling and unifying allosteric activation mechanism in STKR1 kinases that reconciles a large number of experimental studies and sheds light on a novel therapeutic avenue to target disease-related STKR1 mutants.

**AUTHOR SUMMARY:** Kinases play central role in essential physiological process and are attractive therapeutic drug targets. One of the important kinase families is Type 1 Serine/Threonine Kinase Receptors (STKR1), which control gene expression in response to extracellular growth factors. The activities of STKR1 are tightly controlled by their regulatory domain, which is distant from the kinase catalytic site. The underlying molecular mechanism is elucidated here. We identified that formation or disruption of a highly conserved charge-charge interaction located near the ATP binding site, mediates the physiological inhibition or activation of STKR1. We find that the stability of this charge-charge interaction is remotely controlled by interactions propagated from the distant regulatory domain. Several disease-causing mutations are located at the regulatory domain. We demonstrate how those mutations bypass these endogenous STKR1 inhibition mechanisms to produce pathological phenotypes. This study provides a general activation mechanism in STKR1 kinases, thus may benefit understanding the molecular mechanism of diseases and drug development.

## INTRODUCTION

Serine/Threonine Kinase Receptors (STKRs), also known as Transforming Growth Factor *beta* (TGF-β) receptors, are ubiquitous transmembrane proteins that play central roles in many biological processes ranging from cell differentiation, migration, proliferation and adhesion to development ^1,2^. These receptors phosphorylate transcription factor Smad proteins in response to extracellular growth factors including TGF-β, activins, inhibins, nodal, and Bone Morphogenic Proteins (BMPs). In mammals, twelve known STKRs are organized into either type 1 receptors (STKR1) or type 2 receptors (STKR2). STKR1s encompasses seven members named Activin-Like Receptors 1-7 (ALK1–7), while STKR2s consist of five members: Activin Receptor Type 2A, Activin Receptor Type 2B, BMP Receptor Type 2, TGF-β receptor Type 2 and Anti-Mullerian Hormone Receptor Type 2. Extracellular ligands usually promote the formation of a heteromeric complex consisting of two STKR1s and two STKR2s, although in some cases the existence of a pre-formed complex has been reported ^3^. This complex-ligand aggregate enables constitutively active STKR2 ^4–6^ to phosphorylate a glycine-serine (GS) regulatory region in the intracellular domain of STKR1. In turn, STKR2-mediated phosphorylation of the STKR1 GS region enables STKR1 kinase domain to bind adenosine triphosphate (ATP) and transfer its γ-phosphate to downstream Smad substrates. Activated ALK2/3/6 phosphorylate downstream Smad1/5/8 to transduce BMP signaling whereas activated ALK4/5/7 phosphorylate downstream Smad2/3 to transduce TGF-β and activin signaling (Fig. 1*a-b*) ^7^.

**Figure 1:**
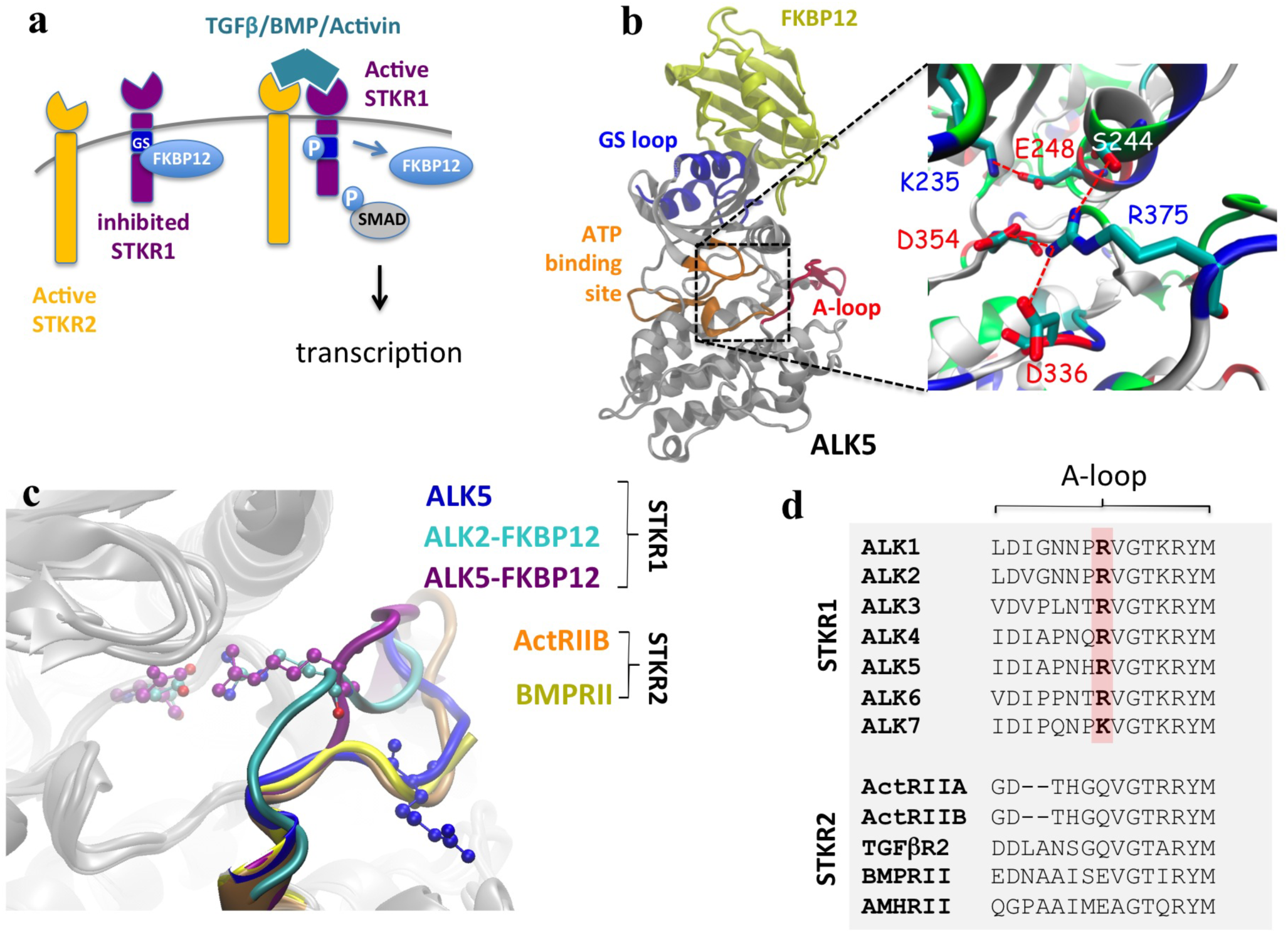
A conserved electrostatic lock in the kinase domain prevents STKR1 activation. (**a**) Scheme of TGFβ/BMP/Activin induced signaling. (**b**) Structure of FKBP12-ALK2 complex and a close view of the R375-D354 salt bridge, as well as other salt bridges and hydrogen bonds involving R375 in ALK2 are illustrated. A conserved Lys-Glu pair (K235-E248) between β3 sheet and αC helix is also shown here. (**c**) Structural alignment of the kinase domain from nine crystal structures. R-D salt bridge residues are shown in CPK mode. Five ALK5 crystal structures without FKBP12 bound are taken from PDB 1IAS, 2WOT, 3TZM, 4X2N, and 5E8S. The overlapped A-loop is shown in blue. ALK2 with FKBP12 bound is shown in cyan (PDB 3H9R) and ALK5 with FKBP12 bound is in purple (PDB 1B6C). For STKR2, ActRIIB is shown in orange (PDB 2QLU) and BMPRII in yellow (PDB 3G2F). The missing A-loop in ALK2 crystal structure (residues 362-374) was transplanted from the crystal structure of inactive ALK2 structure PDB 3Q4U. Small molecular inhibitors in ATP binding site of the crystal structures are not shown. (**d**) Partial sequence alignment of human STKR1 (ALK1-7) and STKR2 isoforms (ActRIIA and B, TGFβR2, BMPRII and AMHRII) showing strict conservation of the positively-charged electrostatic lock in all SKTR1 isoforms while being absent in all constitutively active SKTR2 isoforms.

In the absence of ligand, STKR1 kinase activity is physiologically inhibited by binding of a negative regulator protein FKBP12 to the GS domain. Gain-of-function mutations in the GS and kinase domains are somehow able to bypass FKBP12-mediated inhibition and produce aberrant STKR1 signaling that is associated with various diseases such as heterotopic ossifications and cancer ^8–11^. How STKR1 GS domain modifications by FKBP12 dissociation, STKR2 phosphorylation, or disease-causing mutations lead to activation of the kinase catalytic domain located about 30Å away is unclear. To this aim, structural studies have shown that in the inactive forms of ALK2 and ALK5, ATP binding is prevented by an inhibitory salt-bridge between an aspartate residue located in the conserved DLG kinase motif and an arginine located in the activation loop (A-loop) ^9,12^. However, it remains unknown whether this salt bridge represents a common inhibitory mechanism amongst all STKR1s, and if so, how the salt bridge is allosterically regulated by physiological and pathological modifications of the distant GS domain. In this study, using bioinformatics, molecular dynamics (MD) simulations and experimental assays, we show that this salt bridge acts as a common electrostatic lock among all inactive STKR1s. GS domain modifications allosterically destabilize this electrostatic interaction to induce STKR1 kinase activation. Experimental disruption of this electrostatic lock by mutations of the conserved A-loop arginine in three tested STKR1s produce disease-mimicking leaky signaling in the absence but not presence of the FKBP12 repressor, indicating this electrostatic lock functions as a secondary endogenous repressor downstream of FKBP12.

## RESULTS

### A conserved salt-bridge exists in inactive STKR1 kinase domains

To gain insight into the STKR1 activation mechanism, we aligned nine high-resolution X-ray structures of constitutively active STKR2s (ActRIIB and BMPRII), FKBP12-bound inactive STKR1s and FKBP12-free STKR1s with substrate analogs bound to their catalytic sites. The overall structures of the kinase domains overlap extremely well except for the A-loop (Fig. 1*c*). In the inactive structures, an Arg (R) residue on the A-loop forms an inhibitory salt-bridge with the Asp (D) residue in the conserved DLG motif as previously reported ^9,12^. In contrast, in constitutively active STKR2s and the FKBP12-free ALK5 structures, this A-loop R residue is systematically flipped away from the ATP binding site. Interestingly, a positively-charged residue Arg or Lys at this position is strictly conserved in all STKR1 but not present in any constitutively active STKR2s (Fig. 1*d*). These observations strongly suggest that the R-D salt bridge is ubiquitous in all known STKR1 members and constitutes an electrostatic lock to physiologically prevent STKR1 kinase catalytic function in the absence of ligand.

### FKBP12 dissociation or GS domain phosphorylation destabilizes the electrostatic lock in ALK2

If our hypothesis that the R-D salt bridge acts as an electrostatic lock in STKR1 kinase inhibition is correct, disrupting this electrostatic lock would be necessary during STKR1 kinase activation. To examine this concept, we first carried out multiple MD simulations in explicit solvent to study whether FKBP12 dissociation and GS loop phosphorylation disrupt the electrostatic lock in ALK2. To this aim, four structure systems were prepared from the same crystal structure of inactive FKBP12-ALK2^WT^, namely FKBP12-ALK2^WT^; ALK2^WT^ (no FKBP12); FKBP12-ALK2^WT-Phosp^; and ALK2^WT-Phosp^ (no FKBP12). Phosphorylation sites of ALK2^WT-Phosp^ were selected based on a previous ALK5 study as they are conserved among STKR1 ^7^ (see Methods). The length of each simulation varies between 280 to 300 nanoseconds (ns). In either the absence of FKBP12 or presence of phosphorylation, the R-D lock between D354 and R375 rapidly breaks during the simulation, but remains only stable in FKBP12-ALK2^WT^ simulations (Fig. 2*a-d*). This unique stability of the R-D lock in FKBP12-ALK2^WT^ was double-checked using a duplicated simulation of 280 ns starting from a different snapshot and initial velocity (Fig. S1a). Those simulation results suggest that activation of ALK2 kinase by either FKBP12 dissociation or GS phosphorylation indeed destabilizes the electrostatic lock in ALK2. Interestingly in the simulation of ALK2^WT^ (no FKBP12), the R (Cζ)-D (Cγ) distance varies between ~4 Å and ~8 Å, indicating the salt-bridge reversibly forms and breaks in the absence of FKBP12 (Fig. 2*b*). In contrast, in phosphorylated systems with or without FKBP12, the R-D salt-bridge breaks at around 200 ns and oscillates between ~8 Å and ~14 Å, consistent with the requirement of physiological phosphorylation of GS domain for full activation of ALK2 (Fig. 2c-*d*). We also observed that GS domain phosphorylation results in partial FKBP12 dissociation (Fig. S1b), consistent with previous reports that FKBP12 only binds to unphosphorylated STKR1s ^7,13,14^.

**Figure 2:**
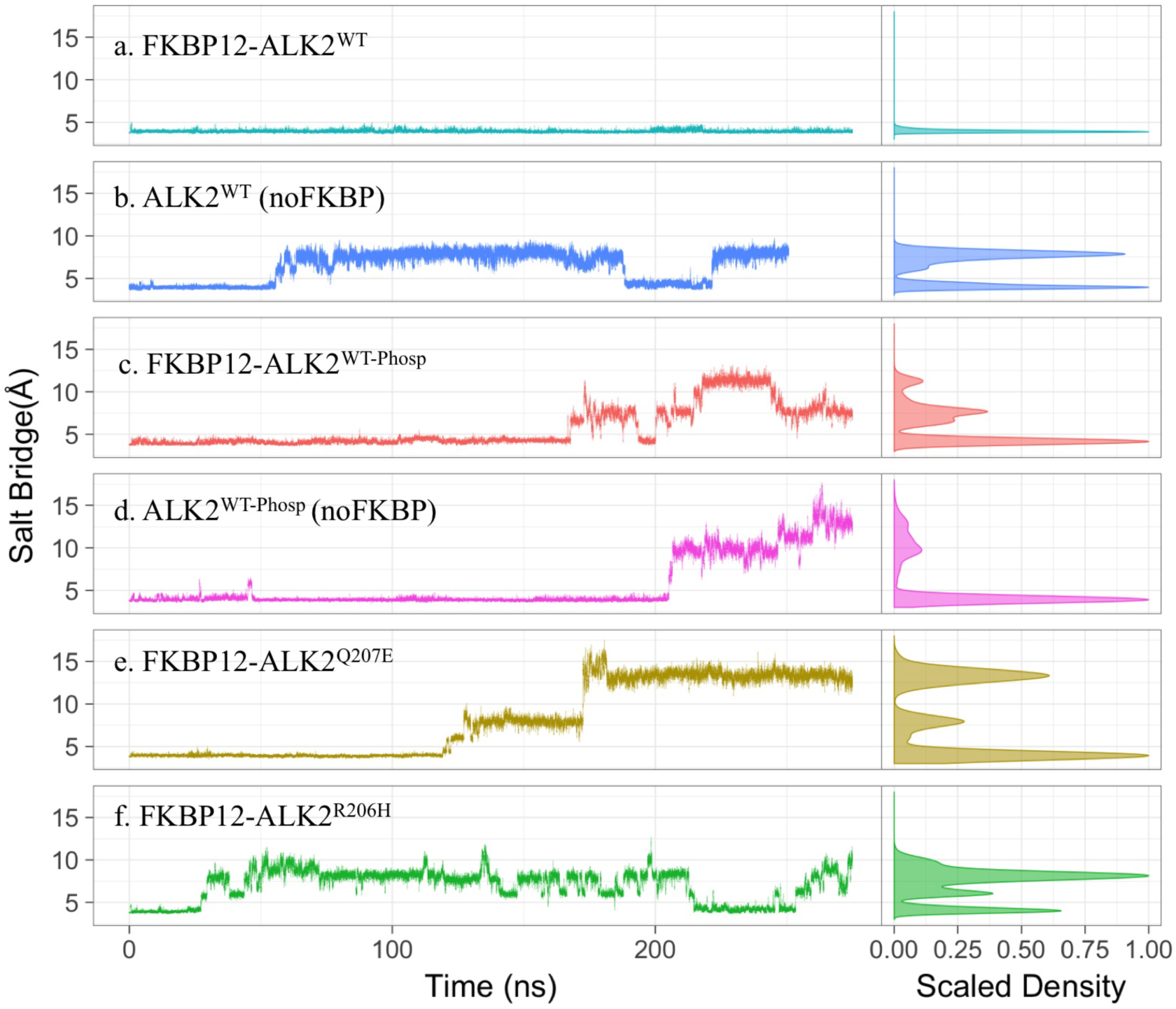
Salt bridge distance between R375(Cζ) and D354 (Cγ) in six simulated systems: FKBP12-ALK2^WT^; ALK2^WT^; FKBP12-ALK2^WT-Phosp^; ALK2^WT-Phosp^; FKBP12-ALK^Q207E^; and FKBP12-ALK2^R206H^. The right panels are the distribution of the distances over the whole trajectories. R375(Cζ)-D354 (Cγ) distance below 5 Å represents salt bridge forming.

### Gain-of-function disease-causing mutations also destabilize the R-D lock in ALK2

The common ALK2^R206H^ and ALK2^Q207E^ gain-of-function mutations in the GS domain produce leaky BMP signaling and are associated with Fibrodysplasia Ossificans Progressiva (FOP) and childhood brainstem tumor diffuse intrinsic pontine glioma (DIPG) ^15,16^. To investigate whether the ALK2 R-D lock is allosterically altered by these mutations, we mutated *in silico* the FKBP12-ALK2^WT^ crystal structure to ALK2^R206H^ and ALK^Q207E^ and performed MD simulations.

Because R206H is partially buried inside the protein structure at the FKBP12 binding interface, its p*K*_a_ (consequently its protonation state) may fluctuate depending on local side chain rearrangements. We, therefore, calculated p*K*_a_ of R206H using constant-pH molecular dynamics simulation ^17^ and PROPKA method (10,000 conformations) ^18,19^. Both constant-pH simulation and PROPKA predict an average p*K*_a_ value of around 6.3, similar to the His p*K*_a_ value in aqueous solvent (Fig. S2, S3). The dominant protonation state of R206H predicted by constant-pH simulation (uncharged state with hydrogen on the Nδ atom) is used for FKBP12- ALK2^R206H^ MD simulation.

It is very interesting that despite the fact that R206H and Q207E are located about 30 Å away from the A-loop, both mutations rapidly destabilize the R-D salt bridge during the 300ns MD simulation (Fig. 2*e-f*). Unlike phosphorylation, R206H and Q207E did not seem to affect FKBP12 binding during the simulations (Fig. S1), and the distribution of the R-D distance in the R206H mutant is similar to the ALK2^WT^ simulation, suggesting that R206H alleviates the inhibitory effect of FKBP12 rather than disrupting its binding to ALK2. This mechanistic hypothesis agrees well with experimental evidence that R206H does not dissociate ALK2/FKBP12 interaction ^9^. In contrast, the R-D distance distribution of the Q207E mutant is similar to the phosphorylation systems, suggesting that the negative charge introduced by Q207E could partly mimic GS domain phosphorylation, potentially leading to partial FKBP12 dissociation in a longer time scale. This mechanism fits well with experimental data that FKBP12 removal does not significantly increase leaky BMP signaling through ALK2^Q207E^ ^9^.

### Flipping of the A-loop arginine corresponds to a 2^nd^ free energy barrier during ALK2 activation

To overcome sampling limitations of the MD simulations, we calculated the free energy profiles or potential of mean force (PMF) along the R-D distance in the ALK2^WT^ system using *Hamiltonian* Replica-Exchange Umbrella Sampling (H-REUS) molecular dynamics simulations ^20–23^ (Fig. 3). The reaction coordinate of the umbrella sampling is the distance between center of mass of hydrogen atoms of one NH2 moiety of R375 guanidine group and one oxygen atom of D354 carboxylic group. A correlation plot for R375-D354 salt bridge distance metrics *vs.* umbrella sampling restraint center reveals clearly that there is a sidechain flipping at 4 Å window and the R-D bridge breaks between 4 Å and 5.5 Å windows (Fig. S7). Three main free energy wells were found at ~4 Å, ~8 Å, and ~14 Å, which are in excellent agreement with the R-D distance distribution from our brute-force MD simulations (Fig. 2a-f). The first free energy barrier in PMF profile is between the energy minima at ~4 Å and ~8 Å, corresponding to the reversible disruption of the R-D salt-bridge, which is seen in the 300 ns equilibrium simulation of ALK2^WT^ without FKBP12 (Fig. 2b) and R206H mutant (Fig. 2f). A higher free energy barrier in PMF profile located at ~10 Å corresponds to R375 passing under the αC helix during flipping out. Only the brute-force simulations of phosphorylated systems (Fig. 2c and d) and Q207E mutant (Fig. 2e) sampled an R-D distance of ~14 Å (Fig. 2), indicating that phosphorylation and Q207E mutation significantly lower a 2^nd^ free energy barrier corresponding to Arg flipping.

**Figure 3:**
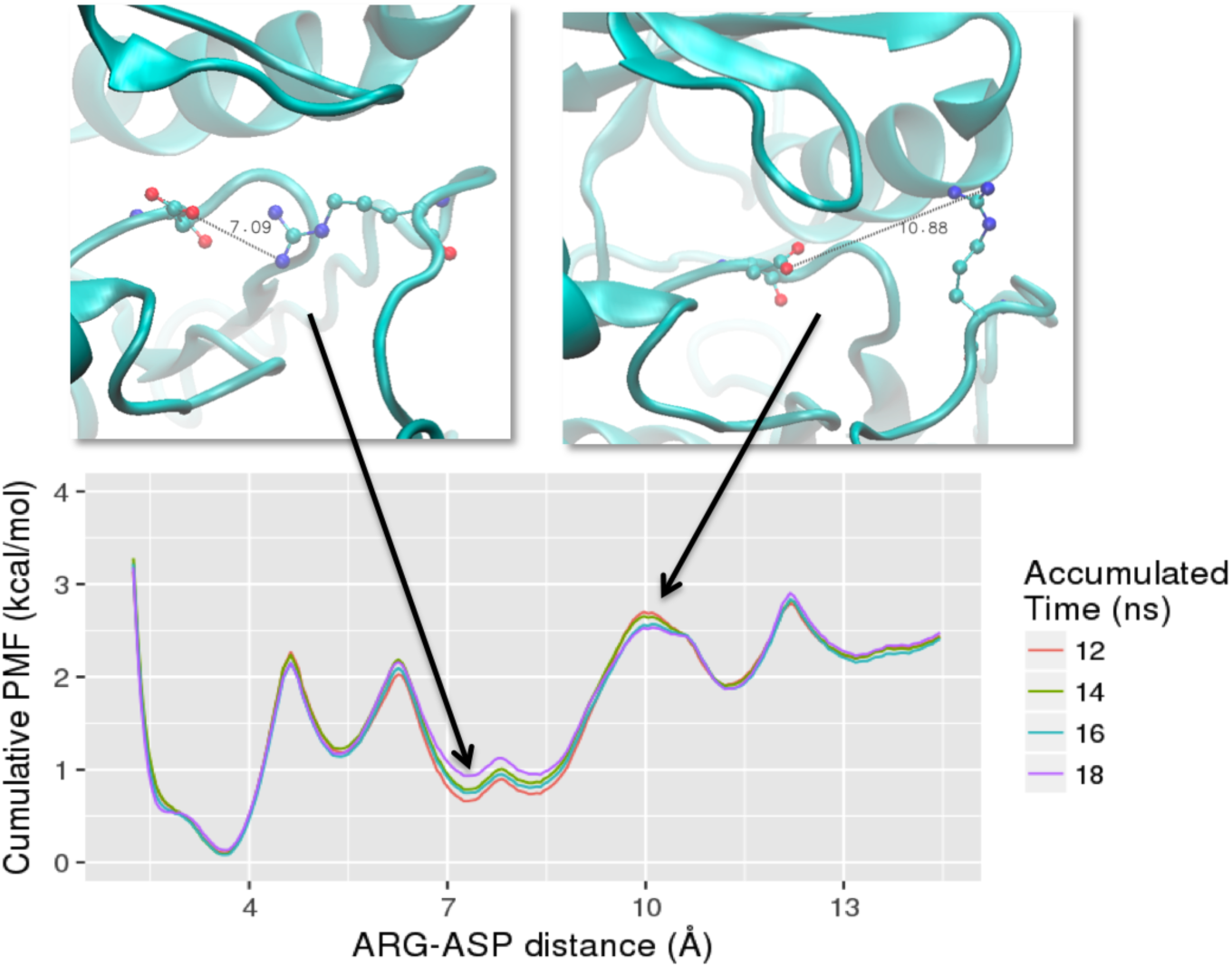
Potential of mean force (PMF) along the R-D distance calculated from *Hamiltonian* replica-exchange umbrella sampling (H-REUS) molecular dynamics simulations. The system used is the ALK2^WT^ and representative snapshots are shown above. A total of 30 ns per window (window width is 1.5 Å) were sampled. To check convergence, PMFs were generated for cumulative windows of the last 18 ns of trajectory. These corresponded to 12-24 ns (labeled as 12 ns in length), 12-26 ns (labeled as 14 ns in length), 12-28 ns (labeled as 16 ns in length), and 12-30 (labeled as 18 ns in length).

### Phosphorylation of the GS loop allosterically regulates the kinase activation in ALK2

To understand how phosphorylation at GS loop allosterically controls the R-D distance at catalytic site, we performed wavelet analysis feature extraction (WAFEX) analysis ^24^ by using the MD trajectories from four systems: FKBP12-ALK2^WT^, ALK2^WT^, FKBP12-ALK2^WT-Phosp^, and ALK2^WT-Phosp^ (Fig. 4). By examining the motion of kinase domain in both physical space (indicated by residue ID on *y*-axis) and temporal space (indicated by simulation time on *x*-axis), we found that ALK2^WT-Phosp^ system undergoes much larger scale motion than FKBP12 dissociation alone (ALK2^WT^) or GS loop phosphorylation with FKBP12 present (FKBP12-ALK2^WT-Phosp^). In addition, WAFEX identified that the most intensive clustered motion (shown in red color) in ALK2^WT-Phosp^ is located at αC helix and β3 sheet. The trajectory frames can be further split into three frame sets based on the discontinuity of the motions shown in the WAFEX diagram (Fig. 4). The jump in the motion of αC helix around 200 ns correlates well with the jump in salt bridge distance in ALK2^WT-Phosp^ (Fig. 2d). Based on this temporal key information, we performed principal component analysis (PCA) using frame set 2 to monitor the change in the motion. PCA revealed that ALK2^WT-Phosp^ system exhibits significantly larger scale motion in GS domain, L45 loop, and αC helix, in comparison with ALK2^WT-FKBP12^ (movie S1). Most interestingly, the direction of motion captured by PCA revealed a separation between αC helix and residue R375 in the R-D lock, as shown by the projections of the R-component vectors (i.e. eigenvectors normalized by their eigenvalues) onto the α carbons (Fig. 5 Top). This separation motion was quantified by calculating natural log of R375 to αC helix separation for first 15 principal components (Fig. 5 Bottom). In frame set 2, the separation is significantly larger in phosphorylated system in the top three principal components, which constitutes 60-75% of the global motion (see cumulative covariance in Fig. S4). Such separation represents an opening motion of the ATP binding site induced by phosphorylation of GS domain, which explains the larger R-D distances observed in the MD simulation of phosphorylated systems.

**Figure 4:**
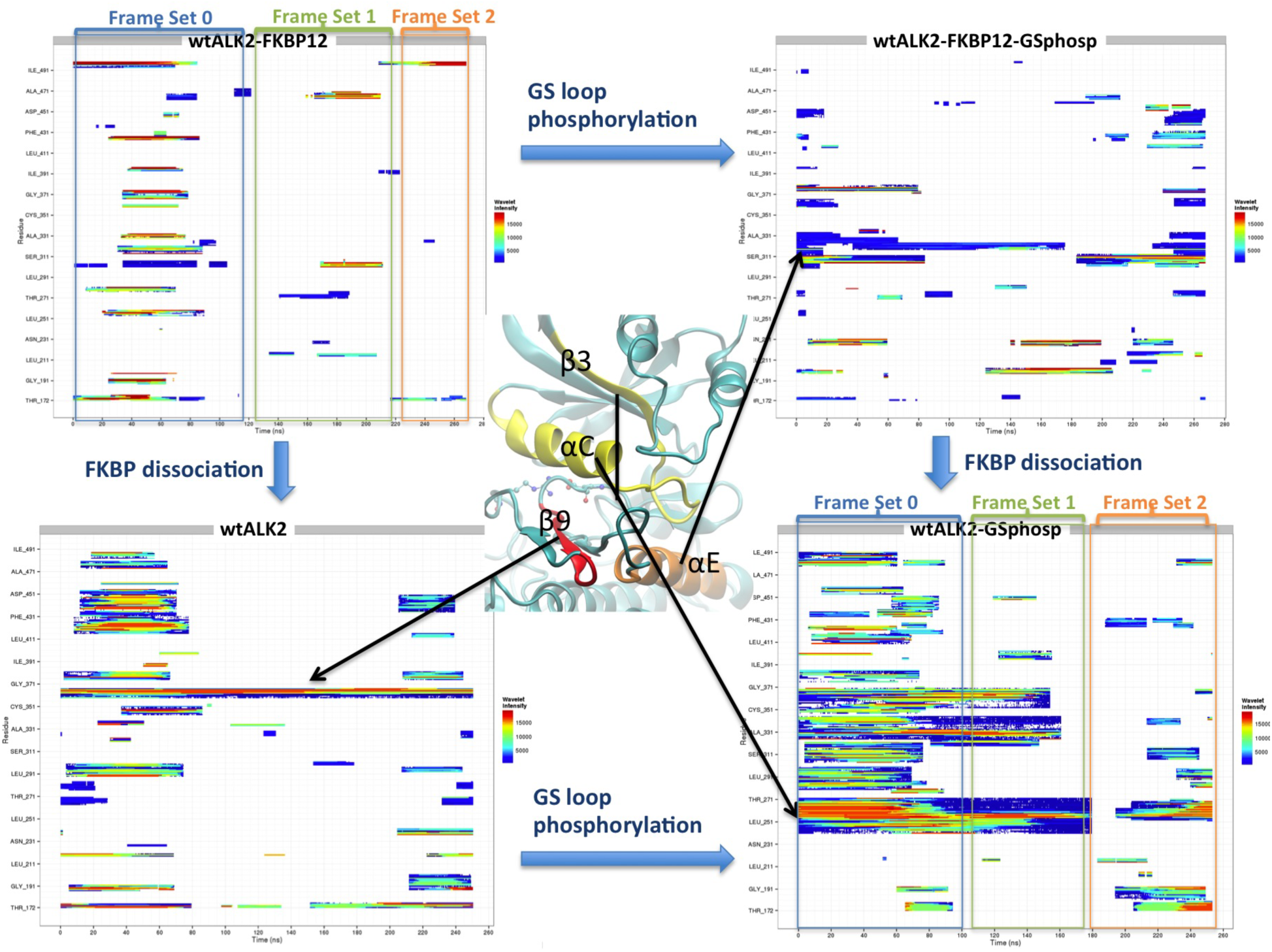
WAFEX (clustered wavelet transform) analysis diagram for trajectories from four systems: FKBP12-ALK2^WT^, ALK2^WT^, FKBP12-ALK2^WT-Phosp^, and ALK2^WT-Phosp^. The simulation time is shown on *x*-axis and residue numbers are shown on *y*-axis. Three-dimensional representation of regions of interest is shown on the left to indicate corresponding regions in the WAFEX plots. Color code indicates the wavelet intensity, with red representing the most intensive and blue the least intensive. Frame sets 0 to 2 were used to guide subsequent principal component analysis.

**Figure 5:**
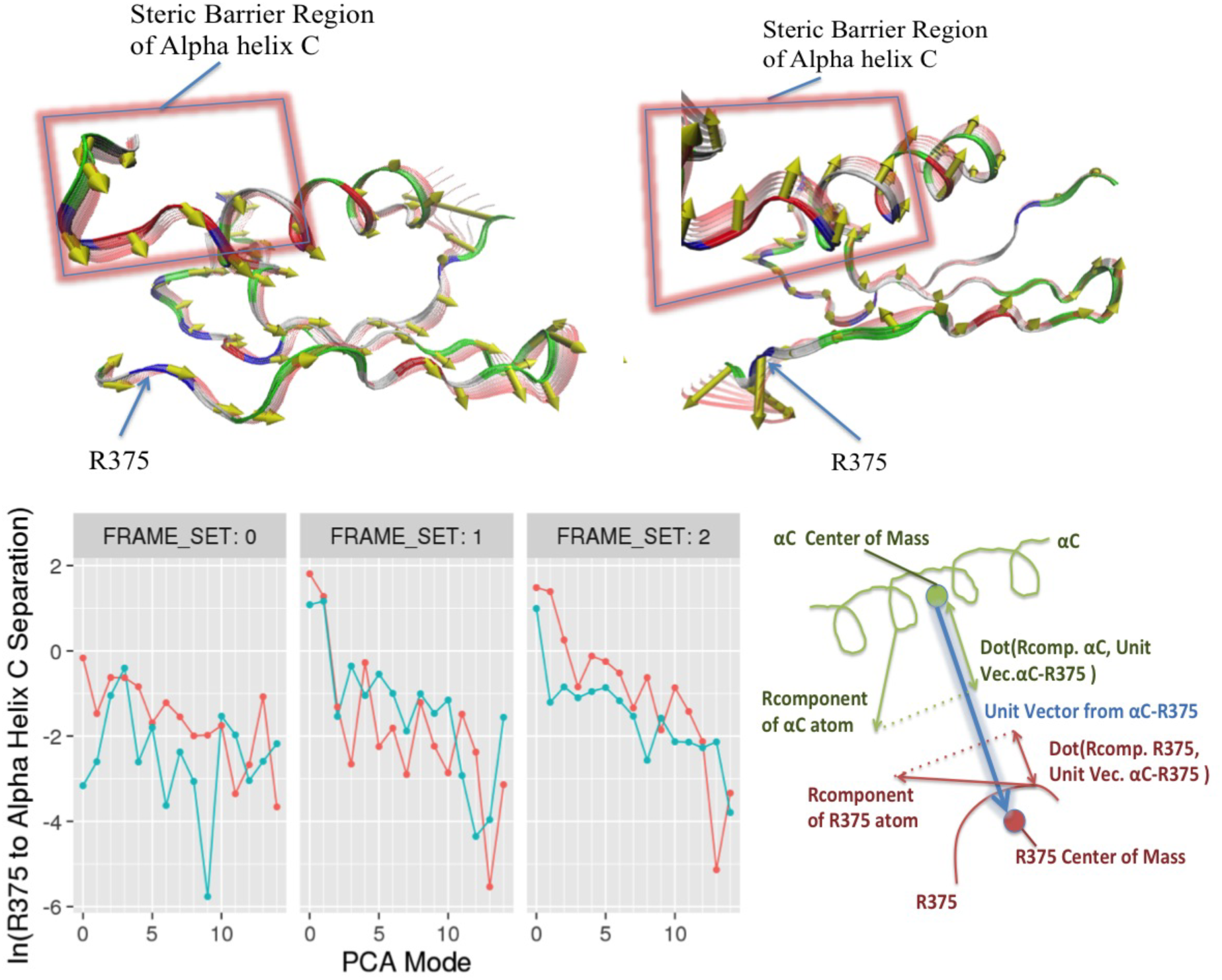
Separation between R375 and αC helix described by principal component analysis. Top. Three-dimensional projections of the second principal component of FKBP12-ALK2^WT^ (left) and ALK2^WT-Phosp^ (right) using frame set 2 trajectories. Projections of the R-Component vectors onto the alpha carbons of each residue are shown as yellow conical tipped arrows. Backbone fluctuation is displayed as red transparent ribbon overlays. The region of αC helix responsible for the steric barrier to Arg flipping is highlighted with red glowing rectangles. **Bottom:** Natural log of R375 to αC helix separation for first 15 principal components of FKBP12-ALK2^WT^ (teal) and ALK2^WT-Phosp^ (orange), taken over trajectory subsets indicated in the previous wavelet analysis figure. The diagram of the R375 and αC helix separation calculation is shown on the right.

### Mutagenesis study confirms the R-D lock disruption as a common mechanism for STKR1 kinase activation

To experimentally verify our MD simulation results, we sought to test the functional consequences of substituting the arginine in the R375-D354 lock of ALK2 to glutamine (ALK2^R375Q^) and glutamate (ALK2^R375E^), two common residues found at the homolog position in STKR2, which are constitutively active (Fig. 1d). In addition, since the FOP-related mutation ALK2^R375P^ was previously characterized ^9^, we have also included this mutation in the study as a control. The ALK2^R375Q^, ALK2^R375E^ and ALK2^R375P^ mutants as well as ALK2^WT^ were expressed into BMP-responsive C2C12/BRE-luc cells respectively, followed by treatments with FKBP12 inhibitor FK506. Compared to cells expressing ALK2^WT^, FK506 treatment alone significantly increased BMP-dependent luciferase activity (RLU=22.67±8.19, n=3, p< 0.01) in cells expressing ALK2^R375P^, which is consistent with a previous report ^9^ (Fig. 6a). Interestingly, the two other mutations ALK2^R375Q^ and ALK2^R375E^ produced comparable luciferase activities to the ALK2^R375P^ (RLU=23.67±3.18, n=3, p< 0.001 for ALK2^R375Q^ and RLU=18±8.50, n=3, p< 0.001 for ALK2^R375E^), in good agreement with R-D salt-bridge in ALK2 acting as an additional repressor beyond FKBP12 to prevent ligand-independent leaky signaling. In the ALK2^WT^ transfected cells, a small, albeit insignificant, increase of luciferase activity with FK506 treatment is consistent with our simulation results that FKBP12 dissociation in ALK^WT^ produces a less stable R-D salt-bridge but still maintains a relative high free energy barrier for Arg flipping, thus allowing the salt bridge to quickly reform following disruption.

**Figure 6:**
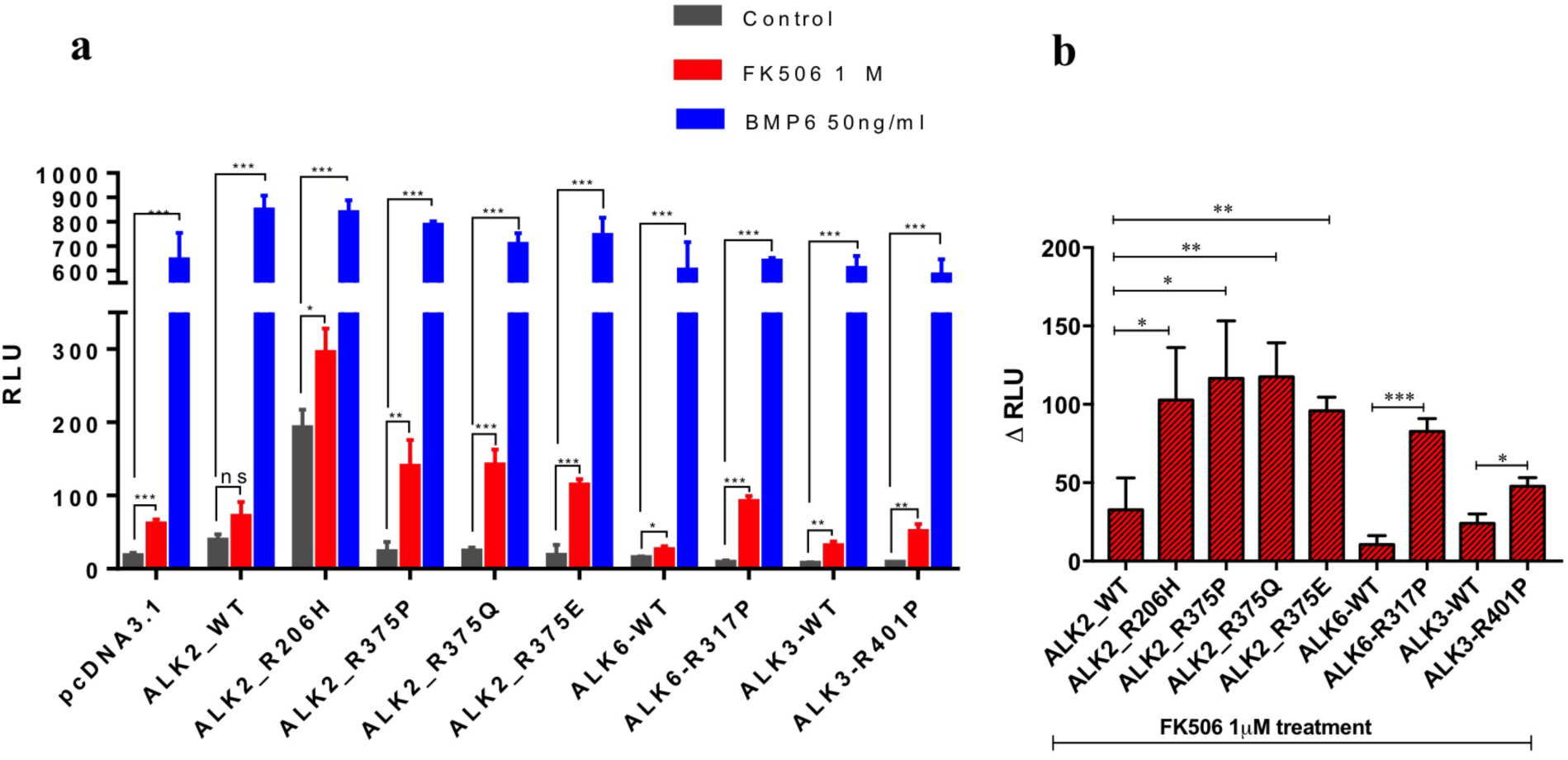
BRE-luciferase reporter assay for different STKR1 receptor mutants. (**a**) C2C12/BRE-luc cells were either untreated (grey bars), treated with 1µM FK506 (red bars) or 50 ng/ml BMP6 (blue bars) overnight, followed by a dual-Luciferase reporter assay. Renilla luciferase reporter pRL-TK plasmid was used as an internal control. (**b**) The relative luciferase unit difference between FK506 treatment and untreatment for the STKR1 receptor mutants. Data are presented as the mean ± S.E.M. (standard error), each performed in triplicate. ns., no significant difference; *p< 0.05, **p< 0.01, ***p< 0.001.

To examine whether an electrostatic lock exists in other BMP type I receptors, we substituted the corresponding arginine in the A-loop of ALK3 and ALK6 to proline as this mutation produces the strong phenotype amongst our tested ALK2 mutations. The ALK3^R401P^ and ALK6^R317P^ mutants were expressed into C2C12/BRE-luc cells respectively, followed by treatments with FK506. Similarly to ALK2 mutants, FK506 treatments dramatically increased luciferase activities in cells transfected with ALK3^R401P^ and ALK6^R317P^ compared to ALK3^WT^ and ALK6^WT^, respectively (Fig. 6*b*). These experiments further demonstrate that the presence of the Arg in the A-loop of ALKs promotes a common inhibitory mechanism independent of the FKBP12 repression. It is noteworthy that the luciferase activities induced by FKBP12 dissociation and Arg mutations are nearly not as high as the ones induced upon addition of BMP6 ligand. This is consistent with previous reports that the phosphorylation on GS domain may facilitate the binding of downstream Smads ^7,25^.

### Network analysis reveals that ALK2^R206H^ allosterically unlock R-D bridge through disruption of FKBP12-ALK2 communication

We noticed that in comparison to the ALK2^WT^ basal luciferase activity (RLU=38.33±4.98, n=3), basal activity (i.e. in the absence of ligand and in the presence of FKBP12) of ALK2^R375P^ (RLU=22.67±8.19, n=3) shows no significant difference, while the basal activity of ALK2^R206H^ is significantly higher (RLU=192±14.50, n=3) (Fig. 6). This is consistent with clinical observations that FOP patients bearing the ALK2^R375P^ mutation generally have milder symptoms compared with patients with the ALK2^R206H^ mutation. Interestingly, R206H is located on the GS domain while the R-D lock is located at the catalytic site 30 Å away. To investigate how ALK2^R206H^ allosterically unlocks the R-D salt bridge, we performed network analysis to determine long-distance correlated residue motions between R206 and R375 in our simulations. Since the geodesic metric network analysis only searches suboptimal paths within a specified cutoff and excludes contributions of contact edges in the entire network, we performed an alternative current-flow betweenness analysis, which provides scoring metric including contributions of all paths (see Methods) ^26–29^. Comparison of ALK2^WT^ versus ALK2^R206H^ mutant networks is shown in Fig. 7. Fig. 7a shows two-dimensional representations of the correlation and current-flow betweenness scores. In the upper plot, the full graph shows that the network of residue contacts involved in transmitting motion between 206 and 375 residues is more focused (i.e. fewer pathways but with bright white or red color) in ALK2^WT^ than ALK2^R206H^. The high transmission sub-networks extracted from the upper panels is shown in the lower panels of Fig. 7a. Interestingly, the contacts between FKBP12 and the kinase domain that are present in ALK2^WT^ (indicated by arrows) disappeared in ALK2^R206H^. To elucidate the relevance of this missing path, the high transmission sub-paths were projected onto the three-dimensional representation of the protein structure. Fig. 7b shows that the missing contacts in the mutant correspond to correlations between T86 of FKBP12 and F246 on ALK2 αC helix. This indicates that the R206H mutation allosterically disrupts FKBP12-mediated stabilization to the R-D lock. Also of note is the edge between P-loop Y219 and R375 show up in ALK2^R206H^, as hydrogen bonding between the Y219 and R375 can be directly observed after R-D lock breaks and R375 enters a metastable intermediate state. In conclusion, the network analysis indicates that the pathological effect of the R206H mutation is to bypass the inhibitory effect of FKBP12 rather than to prevent its binding to the GS domain. This mechanism is consistent with previous co-immunoprecipitation experiments showing that the R206H mutation did not disrupt physical binding of FKBP12 to the GS domain ^9^. Consistent with the previous studies and current luciferase assay, FKBP12 dissociation in ALK2^R206H^ produces a significant increase of luciferase activity as compared to the basal conditions, indicating that R206H does not entirely abolish FKBP12-mediated inhibition.

**Figure 7:**
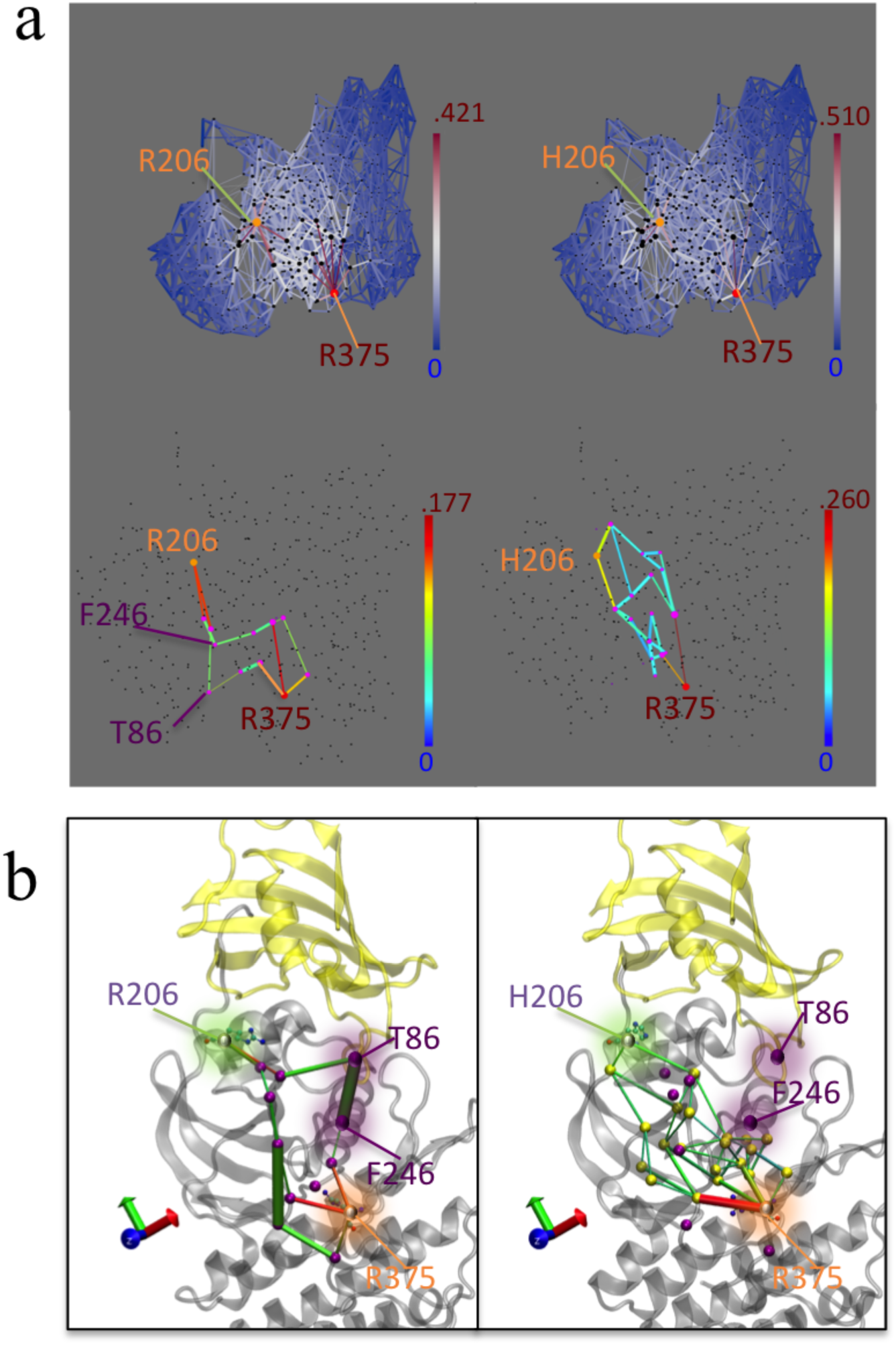
Network analysis shows the communication between R/H206 and R375 in FKBP12-ALK^WT^ (left) and FKBP12-ALK^R206H^ (right). (**a**). 2D depiction of atomic motion transmission networks from residue 206 to residue R375. Edge color depicts current flow-betweenness scores, representing the relative importance of each contact pair with respect to transmission of motion between the residue 206 and R375. Line thickness represents atomic fluctuation correlation between nodes (Cα of residues) Top: full transmission network. Bottom: Sub-networks corresponding to optimum flow-betweenness path plus additional edges of suboptimal paths which exceeded optimal path length by no more than 20%. (**b**) 3D depiction of high transmission networks between residue 206 and R375. Nodes participating in subnetwork, along with the relevant salt bridge lock residue D354 are marked with purple spheres. Nodes present in both wild type and R206H mutant are marked with blue spheres and residue labels while those present in only one of the two networks are marked with purple residue labels. In order to avoid clutter and improve legibility, only nodes present in both R206H and wild type sub-networks were labeled in the R206H sub-network panel.

## DISCUSSION

STKR1 are physiologically activated upon ligand-induced GS domain phosphorylation which promotes dissociation of the FKBP12 repressor, ATP binding to the catalytic site and Smad substrates recruitment. Here we show how the kinase catalytic site becomes active or inhibited in physiological or pathological situations. Modifications in the regulatory GS domain are propagated toward the catalytic domain where they stabilize or destabilize an inhibitory arginine-aspartate salt bridge positioned next to the ATP binding site. Interestingly, this salt bridge is strictly conserved in all STKR1s but is absent in STKR2s which are constitutively active, indicating the requirement of this electrostatic interaction for kinase inhibition.

The salt bridge distance free energy profile reveals three possible states which correspond to the three functional states of the kinase receptors as illustrated in Fig. 8. In absence of ligand and in presence of the FKBP12 repressor, the Asp-Arg salt-bridge is formed and stable, leading to the inhibited form of the STKR1. FKBP12 dissociation and pathogenic mutations in the GS domain lead to transient formation and disruption of the salt-bridge. This state is associated with a leaky or partially active state also called “de-inhibited” state. Finally, the Arg side chain flipping induces a kinase conformation where the salt bridge become permanently disrupted, allowing Asp to participate in ATP coordination. This constitutes the physiological active state of the kinase upon phosphorylation. It is important to notice that experimental removal of the salt bridge alone by mutating Arg did not lead to full kinase activation in a luciferase assay because downstream Smad substrates binding to STKR1 depends upon GS domain phosphorylation ^7,25^. In addition, conformational differences between WT STKR1 and their pathogenic mutants (such as FOP mutant ALK2^R206H^) revealed in our study may offer a novel strategy to design small molecules specific to leaky mutant receptors without interrupt the WT STKR1 activity, thus minimizing potential drug side effects. In summary, our study provides a compelling and unifying general mechanism for STKR1 regulation, and sheds light on a novel therapeutic avenue for targeting disease-related STKR1 mutants.

**Figure 8:**
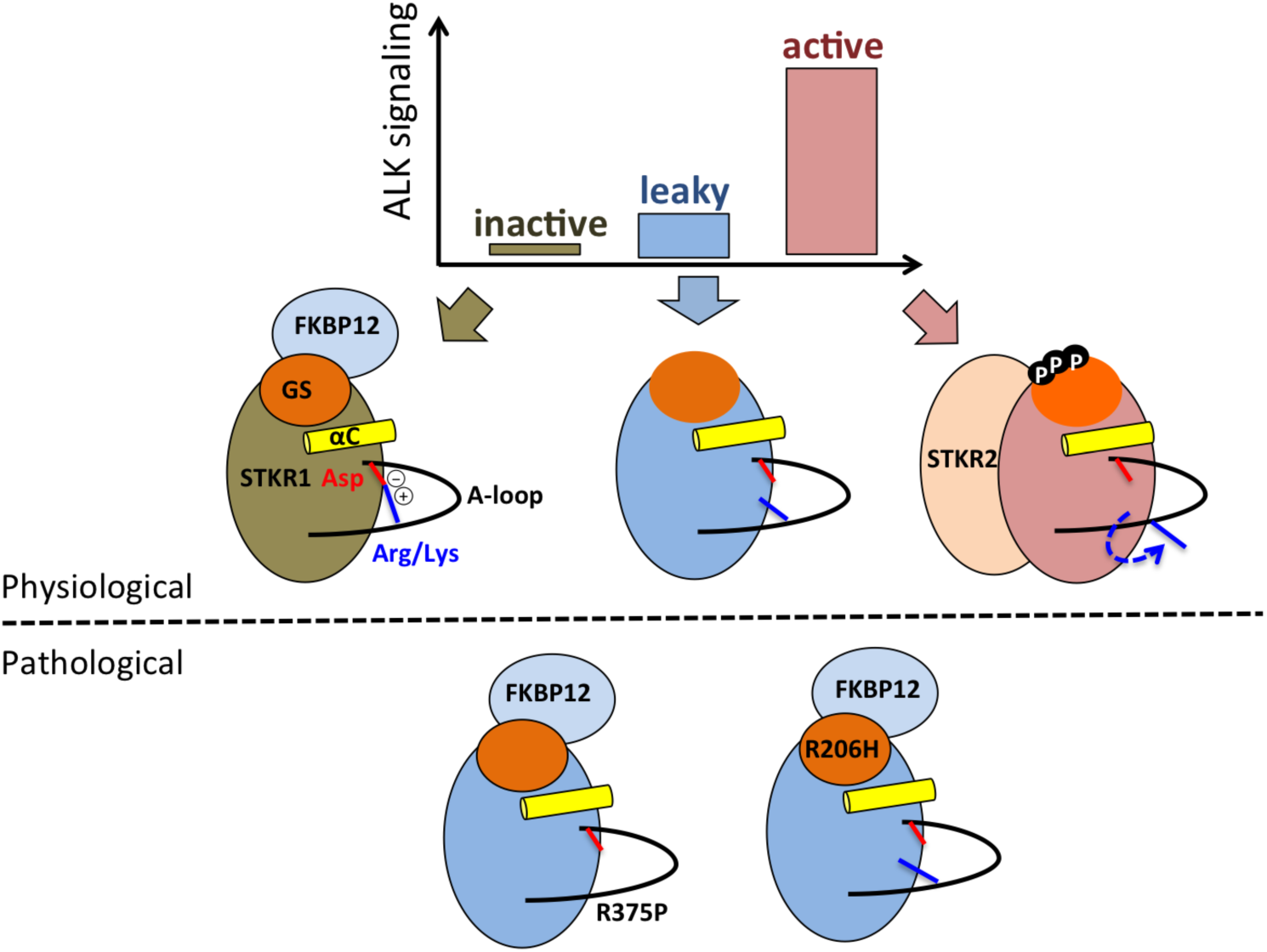
Cartoon showing structure-functions relationships in STKR1s. Various kinase activity levels (inactive, leaky and active) are determined by two inhibitory mechanisms (FKBP12 binding and the endogenous R-D lock) and one activation mechanism (phosphorylation) that promotes FKBP12 dissociation, R-D lock disruption and also promotes substrate binding to the kinase domain.

Five conformational changes are necessary to enable the fully active form of majority of kinases: A-loop opening, DFG-in motion, K-E pair formation, αC helix inward motion, and R-spine formation ^30,31^. Interestingly, all those features already exist in the inactive forms of STKR1 (*i.e.* FKBP12 bound and GS domain unphosphorylated), suggesting that the inactive STKR1 adopts a conformation similar to that of the fully active kinases (Table 1 and Fig. 9a and b). Unlike STKR1s, serine/threonine kinases PKA and Cdk or tyrosine kinases Src or Abl require A-loop phosphorylation for their full activation. Their DFG motifs located at the beginning of the A-loop are tightly coupled to the phosphorylation state of the A-loop. The phosphorylation-induced DFG-in conformation is required for the binding of divalent cations involved in nucleotide recognition, and in forming the regulatory hydrophobic spine (R-spine) in those kinases. In inactive STKR1s, a DLG motif, homolog to the DFG motif, adopts a DFG-in like conformation, but with one main difference that Asp residue is locked by the R-D salt bridge in inactive STKR1 conformation (Fig. 9c). Unlocking this salt bridge enables Asp of DLG motif to align perfectly with the DFG-in conformation of active tyrosine kinases in order to coordinate divalent cations. In summary, we observed that STKR1s adopt an active-like conformation, but are still physiologically inhibited by FKBP12 and the R-D salt bridge. The fact that STKR1 activation requires a much smaller conformational change than other kinases further emphasizes the physiological importance of the R-D lock in STKR1s.

**Table 1.**
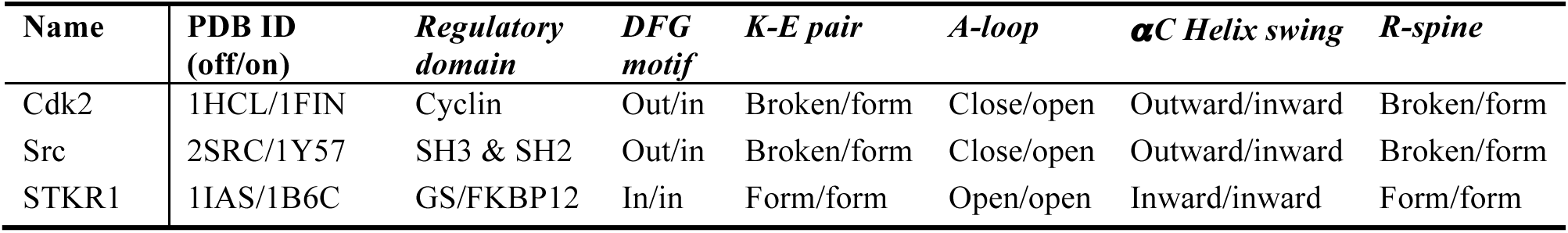
Comparison of key conformational changes from inactive (off) to active (on) state among Cdk2 (cyclin-dependent kinase 2), Src (human tyrosine-protein kinase), and STKT1 ALK5 (TGFβ receptor Type I).

| Name | PDB ID (off/on) | Regulatory domain | DFG motif | K-E pair | A-loop | αC Helix swing | R-spine |
| --- | --- | --- | --- | --- | --- | --- | --- |
| Cdk2 | 1HCL/1FIN | Cyclin | Out/in | Broken/form | Close/open | Outward/inward | Broken/form |
| Src | 2SRC/1Y57 | SH3 & SH2 | Out/in | Broken/form | Close/open | Outward/inward | Broken/form |
| STKR1 | 1IAS/1B6C | GS/FKBP12 | In/in | Form/form | Open/open | Inward/inward | Form/form |

**Figure 9:**
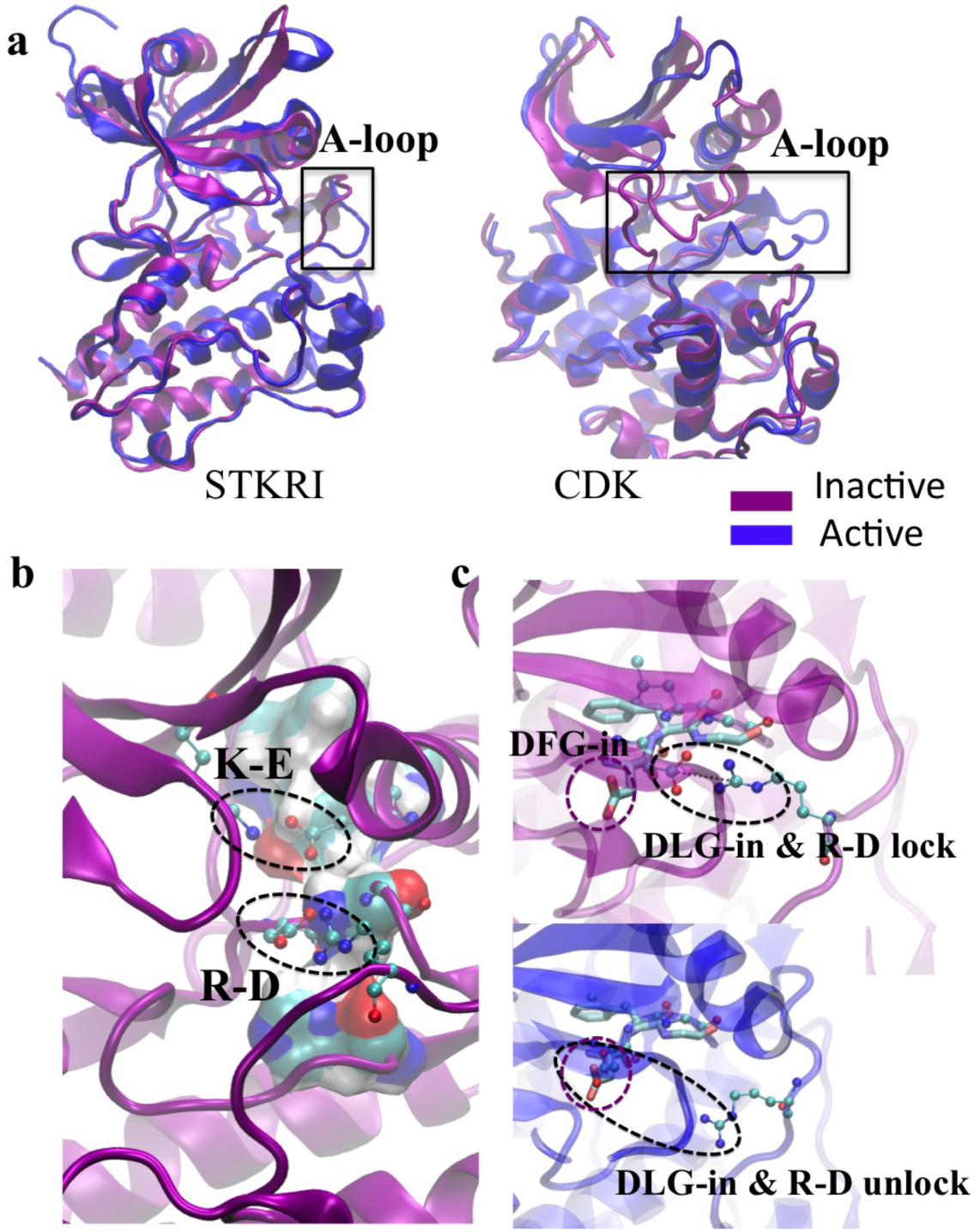
Inactive STKR1 adopts a conformation similar to that of the fully active kinases. (**a**). Comparison between active and inactive A-loop conformation between STKR1 (ALK5), cyclin-dependent kinase (CDK). (**b**). Representative inactive FKBP12-ALK2^WT^ catalytic site conformation from MD simulations, with K-E and R-D salt bridge highlighted. R-spine is shown in vdW surface mode colored by atom type. (**c**). Comparison of DLG-in with DFG-in conformation. DFG motif from Src is shown in licorice mode and GLG motif and R375 are shown in CPK mode.

## METHODS

### Computational programs

All simulations were done using NAMD2.9b ^32^ or AMBER16 ^33^ with CHARMM C36 force field ^34^. WAFEX analysis is done using CPPTRAJ ^35^ and R ^36^. PCA is done using pytraj and visualized via the Normal Mode Wizard plugin of the VMD program ^37,38^. Constant pH simulations were conducted using the constant pH functionality with Generalized Born implicit solvation under the AMBER16. In network analysis, the flow betweenness scores and corresponding networks were computed R script and NetworkView plugin in VMD ^28^.

### System preparation

The initial structure of wtALK2 was taken from the crystal structure of the ALK2-dorsomorphine in complex with FKBP12 (PDB ID: 3H9R) of *Homo sapiens* species. Using VMD, small molecular inhibitor was removed from the crystal structure to eliminate its inhibitory effect on ALK2, and test the inhibitory effect of FKBP12 alone. In this crystal structure, ALK2 partial A-loop (residues 362 to 374), and the β-turn between β4 and β5 (residues 273 to 275) were missing and therefore transplanted from PDB ID 3Q4U. The pKa values were calculated using PROPKA. CHARMM-GUI ^39^ was used to read in the PDB file and generate solvated systems. The phosphorylation site in ALK2 GS domain was selected based on experimental data of ALK5, which defines the exact amino acids of the GS domain (Thr185, Ser187, Ser189 and Ser191) that are phosphorylated during activation ^7^. Each system was solvated in a rectangular water box with 12 Å edge distance from protein surface. Each system was neutralized with K^+^ and Cl^−^ ions at a physiological salt concentration of 150 mM. All simulations employed the all-atom CHARMM C36 force field for proteins and ions, and the CHARMM TIP3P force field ^40^ for water.

### Simulation protocol

All brute-force simulations were performed with NAMD2.9b using periodic boundary conditions at constant temperature and pressure (NPT ensemble) of 300 K and 1 atm using Langevin thermostat and Andersen-Hoover barostat. Long-range electrostatic interactions were treated using the particle-mesh Ewald (PME) method. A smoothing function is applied to both electrostatic and van der Waals forces between 10 Å and 12 Å. The dynamics were propagated using Langevin dynamics with langevin damping coefficient of 1 ps^−1^ and a time step of 2 fs. The non-bonded interaction list was updated on every integration step using a cutoff of 13.5 Å. The SHAKE algorithm was applied to all hydrogen atoms. The molecular dynamics equilibrium was set to relax the atomic system by releasing the harmonic constraints (force constant 50 kcal/mol/Å^2^) stepwise (every 200 ps) on water and ion molecules, protein side chains, and eventually the protein backbone.

### Constant-pH simulation

To assess the protonation state of H206 in the R206H mutant and any consequential perturbation of the surrounding electrostatic environment incurred by this mutation, constant pH simulations were conducted using the constant pH (CPH) functionality of the GPU accelerated PMEMD program with Generalized Born implicit solvation ^41,42^ as available under the AMBER16 modeling and simulation suite. Prior to constant pH simulation, the PDB of the FKBP12-ALK2^R206H^ mutant was preprocessed for use in CPH simulations under AMBER. The hydrogen atoms of the appropriate HIS residue were stripped and the residue renamed to HIP in order to denote it as a titratable HIS residue. The system was then processed for use under constant pH simulation as described in the AMBER16 constant pH tutorial and the corresponding paper by Mongan et. al. ^17^. The system was minimized, then heated and equilibrated for a total of 6 ns, followed by 100 ns of production simulation. Protonation state data for H206 was then extraction from the CPH production simulation using the cphstats tool and imported into R for further analysis.

### Free energy calculation

The energetics of R-D lock dissociation was investigated using Hamiltonian replica exchange umbrella sampling (H-REUS) simulations. The last snapshot of ALK2^WT^ system trajectories was used as a starting point for the HREUS-1D simulations. A set of 1.5 Å width windows was constructed using a harmonic restraint to enforce R-D separation distances. Prior to running HREUS simulation, each window was allowed to equilibrate for a total of 10 ns at its specified separation distance. This was accomplished by iteratively increasing the restraint distance so that each window was equilibrated starting from the endpoint of the previous window’s equilibration trajectory. This was done to minimize distortions generated during rapid separation of the R-D lock. An exchange rate of 0.2 picoseconds (ps) was employed. System energies were output after each exchange event, and separation distances were recorded every 0.02 ps. Trajectory frames were written every 100 ps. The total simulation time is 30ns per window. All simulations were performed using the PMEMD module of the AMBER modeling and simulation package with support for MPI multi-process control and GPU acceleration code. All simulation parameters were implemented to match the standard MD simulations performed in NAMD described above, except the barostat method and VDW cutoff parameter. The standard AMBER cutoff 10 Å radius was used rather than the NAMD implementation which utilizes a smooth function between 8 and 12 Å, in order to avoid significant slowdown in GPU code algorithms. The AMBER simulations used a Monte-Carlo barostat rather than the standard langevin barostat employed for NAMD simulations. This again was necessary to ensure optimal performance of the GPU accelerated simulation code. Post processing was performed on the in-house computing cluster facilities. The PMF was generated utilizing Weighted Histogram Analysis Method (WHAM) ^43^. The PMF data output by WHAM for each sub-trajectory was collected and analyzed using R. Cumulative PMFs over every 2 ns were then plotted to ensure convergence of the PMF at the region of interest. The last 4 cumulative PMF are shown. Only data for the first 9 windows are displayed since the PMF is apparently still converging at much higher separations.

### Wavelet Analysis

The recent paper by Heidari et. al ^24^ describes a novel WAFEX analysis method for elucidation of the temporal and spacial location of relevant motions within a trajectory. Prior to performing WAFEX analysis, the trajectories from the NAMD based simulations were stripped of all solvent molecules and only the heavy atoms of the remaining protein were retained. Further, after a preliminary run, it was noted that torsional motion between the FKBP and kinase domain dominated the wavelet output and obscured other relevant motions of the kinase domain when the binding protein was present and included. Thus, only the kinase domain was considered in this analysis, with the binding protein being stripped from the trajectory prior to conducting WAFEX. A total of 60 timescales ranging from 10 ps (trajectory time step), up to 300 ns (approximate trajectory duration) were used. The Morlet wavelet was employed, necessitating a correction value of 1.01 as described in the paper by Heidri et. al ^24^. The default chi squared cutoff of 1.6094 was used for the noise reduction threshold. A minimum cluster size of 350 points with an epsilon value of 20.0 was specified for the clustering algorithm. The WAFEX analysis along with preliminary preprocessing of trajectories was conducted using the modified CPPTRAJ program. The wavelet and clustering data produced by CPPTRAJ was then processed in R in order to merge the wavelet intensity data with the clusters predicted by the clustering analysis and produce corresponding clustered intensity plots. Silhouette scoring plot for temporal clustering of WAFEX data was generated by computing silhouette scores for consecutive k-cluster cuts taken from the hierarchical clustering of the frame-wise WAFEX data (Fig S5). Hierarchical clustering was computed using Euclidean distance norms over the log of the wavelet intensities at each frame along with Ward similarity scoring as implemented in the hclust function of the fastcluster package in R. The consistency of temporal clustering of WAFEX data by eye and by automated hierarchical clustering cut to 3 clusters is shown in Fig. S6.

### Principal Component Analysis

Since previous analyses indicate that the trajectories produced are likely exhibiting dynamic transition events, it was important to perform principal component analysis (PCA) over relevant subsets of each trajectory in order to ensure that the motions predicted by PCA correspond to the relevant conformation changes occurring in each individual event. For FKBP12-ALK2^WT^ system, these sets were determined to be the times ranging from 0 to 120 ns, 120 to 220 ns and 220 to 250 ns. For ALK2^WT-Phosp^ systems the time ranges were chosen to be 0 to 100 ns, 100 to 180 ns, and 180 to 250 ns. These sub-trajectories were loaded using pytraj (a python implementation of CPPTRAJ and PTRAJ) and the coordinates of backbone atoms extracted by stripping all other atoms from the trajectories. As with wavelet analysis, the FKBP binding domain was stripped as well. Additionally, the first and last 16 residues of the kinase domain were stripped since these regions corresponded to unstructured tails, which exhibited whip-like motions that would dominate the principal components and obscure more relevant motions. The pca function of the pytraj library was then used to perform principal component analysis on each trajectory subset. The resulting eigenvalue and eigenvector data was then exported to data tables for further analysis in R. Additionally, the PCA data for each trajectory subset was output as NMD format files for visualization via the Normal Mode Wizard plugin of the VMD program.

The extent to which each principal component contributed to separation between the R375 residue of the R-D lock and the portion of the αC helix acting as a steric barrier to Arg flipping was then quantified from the PCA data output by pytraj using R. To do so, the unit vector pointing from the center of mass of the steric barrier region of αC toward the center of mass of R375 was computed for the averaged structure generated over each trajectory subset. The average of the dot products of this vector with the R-component vectors ^44^ of each atom of αC was computed as well as the average of the dot products of the separation unit vector with the R375 atom R-component vectors. These two averages were then subtracted to yield the net separation motion described by each principal component. Finally, the Normal Mode Wizard plugin of the VMD visualization program was used to load the NMD format output files generated from the first 15 PCA modes computed using pytraj to produce renderings of representative principal component modes in the R375 / αC region to facilitate visual comparison of the relative motions of R375 and αC helix.

### Network Analysis

Current-flow betweenness provides an alternative scoring metric ^27,29^. This metric starts by constructing a ‘correlation resistance’ network by taking the reciprocal of the correlation for each pair of edge in the contact correlation network. The graph laplacian of this network is then constructed. This corresponds to the matrix constructed such that for any distinct pair of residues in the correlation resistance network, [i,j], the corresponding off diagonal entry of the graph laplacian matrix a_i,j_ is the inverse of the value of the edge in the resistance network. The inverse of this graph laplacian can then be used to compute a ranking for the relative importance of a given contact edge [i,j] in the correlation network, with respect to the transmission of motion between a given set of source residues S and target residues T. The following equation describes the procedure for computing the effective flow betweenness score for a given contact pair (i,j) given the appropriate pseudo inverse matrix:

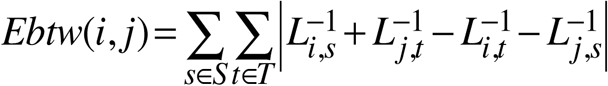

where L^−1^ is the pseudo inverse of the graph laplacian of the correlation resistance network and S and T are the sets of source and target nodes between which motion transfer is being considered. It should be noted that this equation assumes that sets of source and target nodes, S and T, are distinct. If not, the summation must be updated to include only distinct combinations of pairs. Since Laplacian matrices constructed from such graphs are, in general, not directly invertible, the Moore-Penrose pseudo-inverse is used as the effective inverse of the correlation resistance network’s graph laplacian. While the Bozzo and Franceschet paper ^26^ also discusses methods for approximating the pseudo inverse for large methods, direct computation was used here, since our network is relatively small (under 500 nodes).

In brief, this method of computing betweenness scores is equivalent to modeling the correlation network as if it were a network of electrical resistors, wherein each edge is represented as a resistor with resistances equal to the reciprocal of the absolute value of that edge’s correlation. The pseudo inverse of the graph laplacian for this network will then represent the relative difference in electrical potential between each pair of nodes. By selecting a set of nodes as the ‘sources’ and another set as the targets, or ‘grounds’, it is then possible to predict the corresponding ‘relative flow’ across each edge as if 1 unit of ‘current’ (e.g. atomic motion) were sent from the source(s) to the target(s). This ‘relative electrical flow’ can serve as a scoring function (betweenness) to rank each edge or node in order of its relative importance with respect to transmission of motion between the source and target nodes. These betweenness scores can then be used to isolate the most relevant sub-networks that transmit motion between either the FKBP binding interface or residue 206, to residue R375 of the R-D saltbridge lock. The betweenness scores and corresponding optimal sub-networks were computed from the correlation and contact map matrices generated by the NetworkView plugin using R and the corresponding igraph ^45^ R library, and then visualized in two-dimensional format in R and three dimensional format in VMD.

### Molecular biology and mutagenesis

pCDNA3 plasmids harboring ALK2^WT^, ALK3^WT^ and ALK6^WT^ were purchased from Addgene (#80870, #80873 and #80882). Point mutations in ALK2, ALK3 and ALK6 were made in-house using the Q5-Site Directed Mutagenesis Kit from New England Biolabs. Mutagenic oligonucleotides were designed using the NEBaseChanger online tool (New England Biolabs) and purchased from Integrated DNA Technologies. Mutations were confirmed by automated Sanger sequencing (Genewiz).

### Transfection and Luciferase reporter assay

Transient transfection was performed using Fugene HD transfection reagent (Promega) according to manufacturer’s instructions. Briefly, 7.5 × 10^3^ C2C12/BRE-luc cells (stable mouse C2C12 cells expressing the Id1 promoter-firefly luciferase reporter) were seeded in 96-well plates to culture in the growth medium DMEM/10% FBS without antibiotics. After overnight culture, the cells were transfected with pcDNA3.1(+), human wild type ALK2, ALK2^R206H^, ALK2^R375P^, ALK2^R375Q^, ALK2^R375E^, wild type ALK6, ALK6^R317P^, wild type ALK3 and ALK3^R401P^ respectively, along with the Renilla luciferase reporter pRL-TK plasmid. Five hours after transfection, cells were starved in DMEM containing 0.5% FBS for 4 h. Then the cells were untreated or treated with 1µM FK506 (Cayman) or 50 ng/ml BMP6 (R&D) overnight before lysis. Luciferase activities were determined according to the Dual-Luciferase® reporter assay system (Promega) using Renilla for normalization of transfection efficiency. Data are presented as the mean ± S.E.M (standard error), each performed in triplicate. Student’s unpaired t-test was used for two groups of statistical analysis. A p-value <0.05 was considered statistically significant.

#### Code availability

The R scripts used to generate the current flow betweenness scores and suboptimal networks can be found on GitHub at: https://github.com/LynaLuo-Lab/network_analysis_scripts.

## ACKNOWLEDGEMENT

We thank Dr. Aristidis Moustakas for making ALK3 and ALK6 plasmids available through Addgene. Computational work reported here was supported by the National Science Foundation grant number MCB160119 and TG-MCB150042 through the Extreme Science and Engineering Discovery Environment (XSEDE) program. Funding support is from seed funds from Western University of Health Sciences (J.J.L; J.H. and Y.L) and Chinese American Faculty Association (CAFA) faculty development grant (Y.L.).

## AUTHOR CONTRIBUTION

Y.L., J.H., J.J.L., W.M.B-S., and A.A. designed research; W.M.B-S., A.A., P.C., C.X., J.L., J.H. and Y.L. performed research; W.M.B-S., A.A., P.C., C.X., J.L., J.H. and Y.L. analyzed data; Y.L., J.H., J.J.L., W.M.B-S., and A.A. wrote the paper.

## Supporting Information

**Figure S1.**
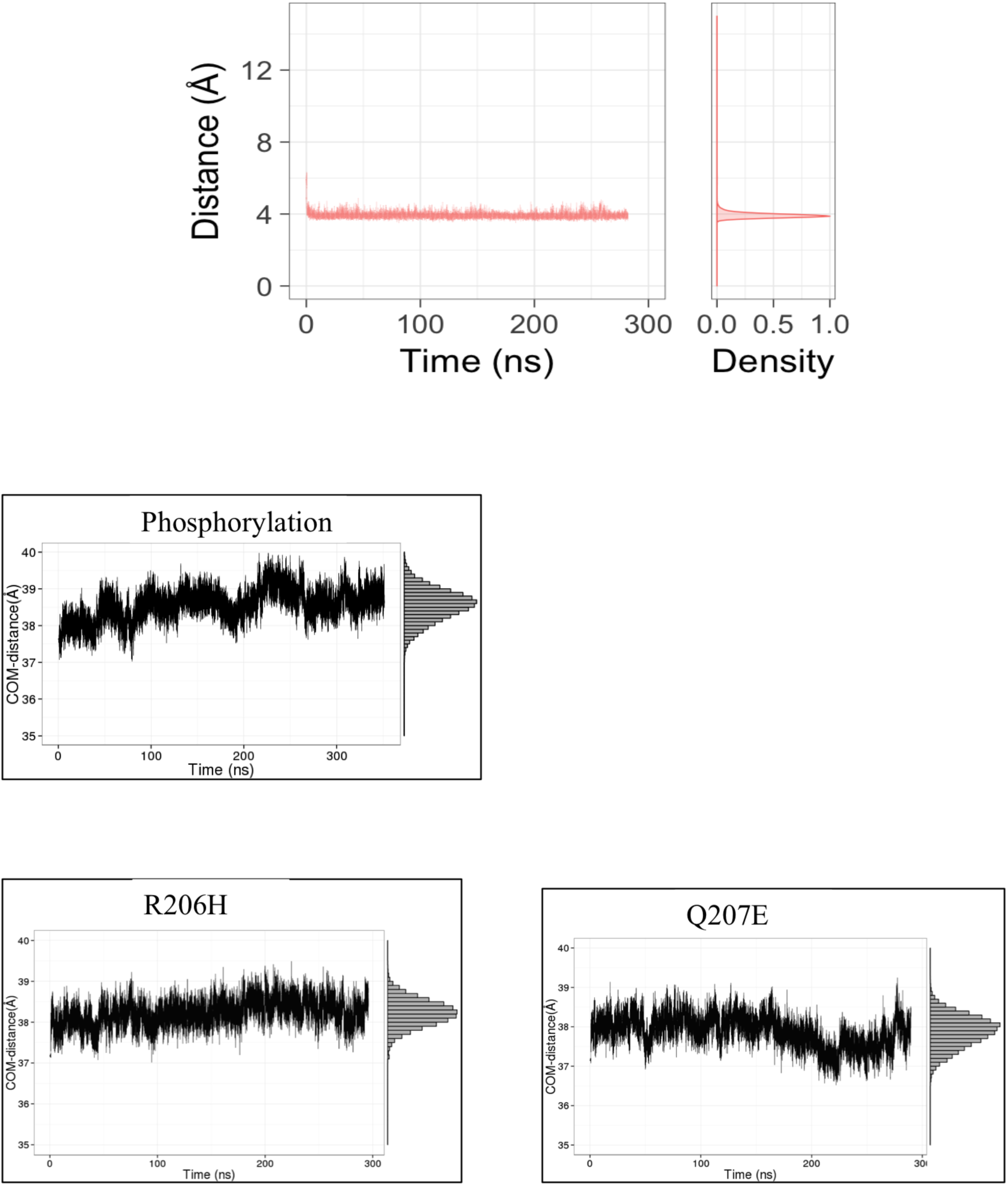
(a) Salt bridge distance between R375(Cζ) and D354 (Cγ) from a duplicated simulation of FKBP12-ALK2^WT^. (b). Center of mass distance between FKBP12 and ALK2 during 300 ns simulations of FKBP12-ALK2^WT-Phosp^, FKBP12-ALK2^R206H^, and FKBP12-ALK2^Q207E^.

**Figure S2.**
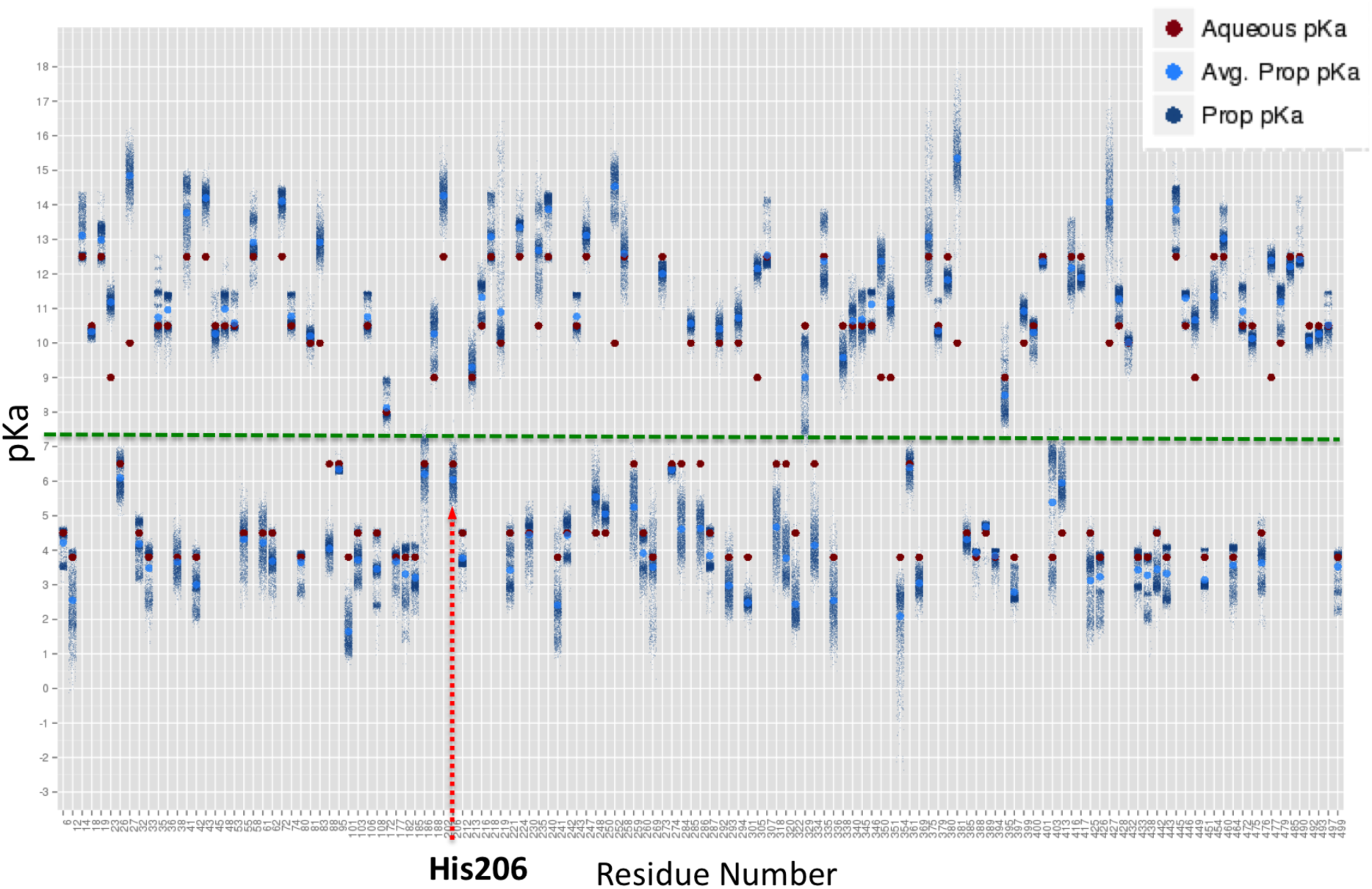
The p*K*a of each residue on ALK^R206H^ calculated by PROPKA using 10,000 snapshots from 300 ns MD simulation trajectory. Dark blue dots represent the p*K*a value from each snapshot, light blue dots represent average value, and red dots represent p*K*a in aqueous solution. Green dashed line indicates physiological pH 7.4. His206 is indicated by red dashed arrow.

**Figure S3.**
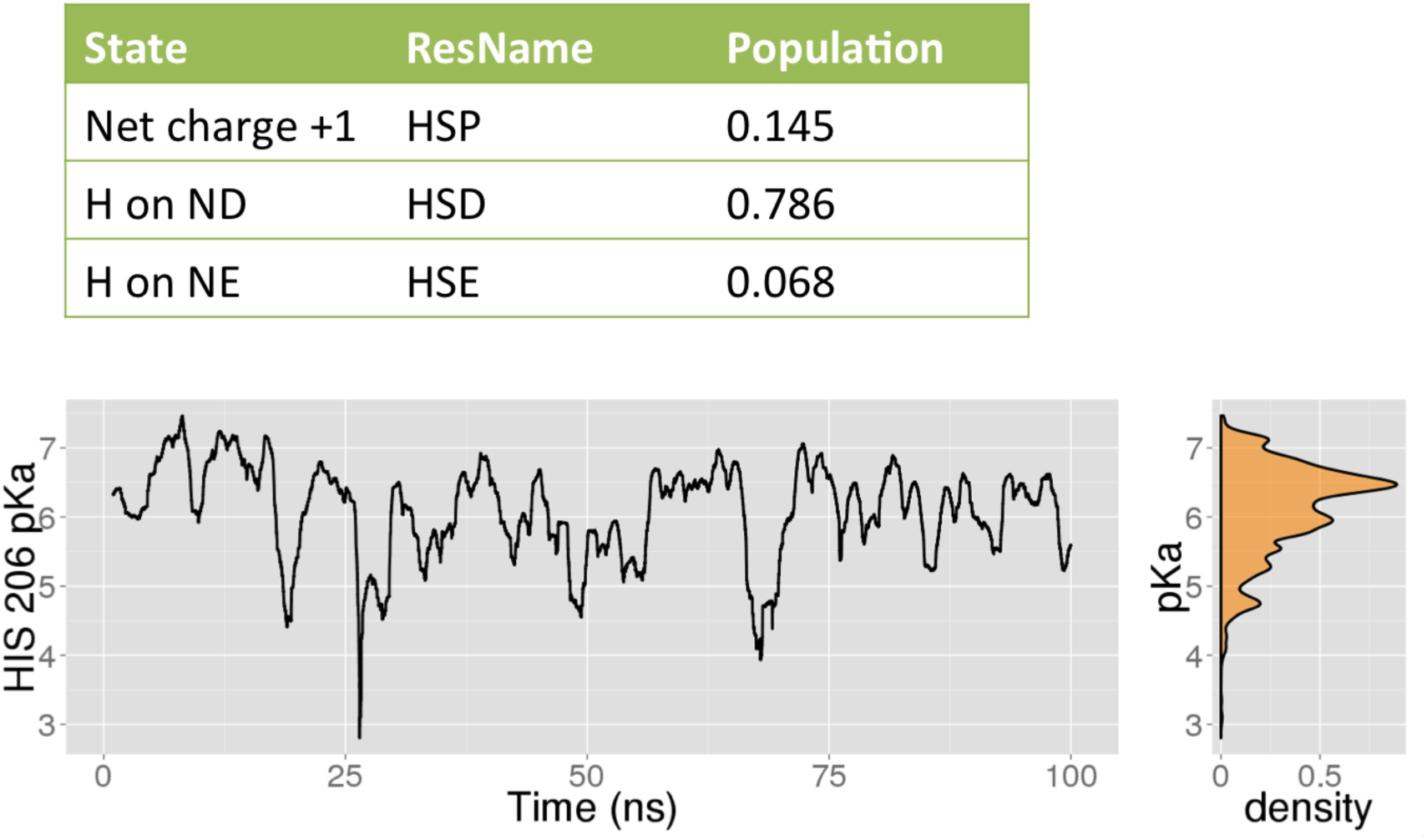
Constant-pH molecular dynamics simulation results. The population of His206 protonation state is shown in the top table. The calculated p*K*a of His206 is plotted against simulation time. Right: a density plot of calculated pKa of His206 during the simulation.

**Figure S4.**
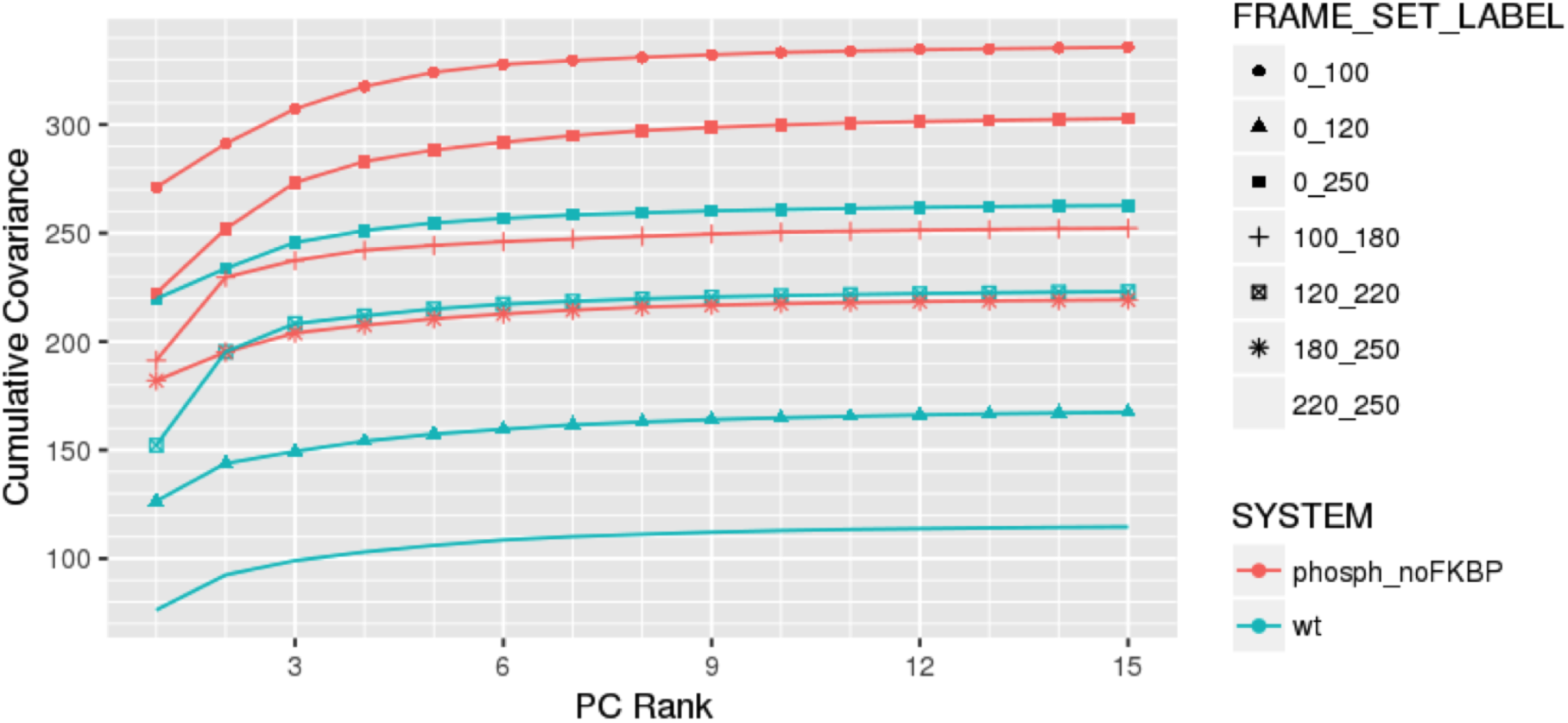
Convergence of cumulative covariance described by principal component modes. FKBP12-ALK2^WT^ is shown in teal and ALK2^WT-Phosp^ is shown in orange, taken over trajectory subsets indicated in the previous wavelet analysis figure.

**Figure S5.**
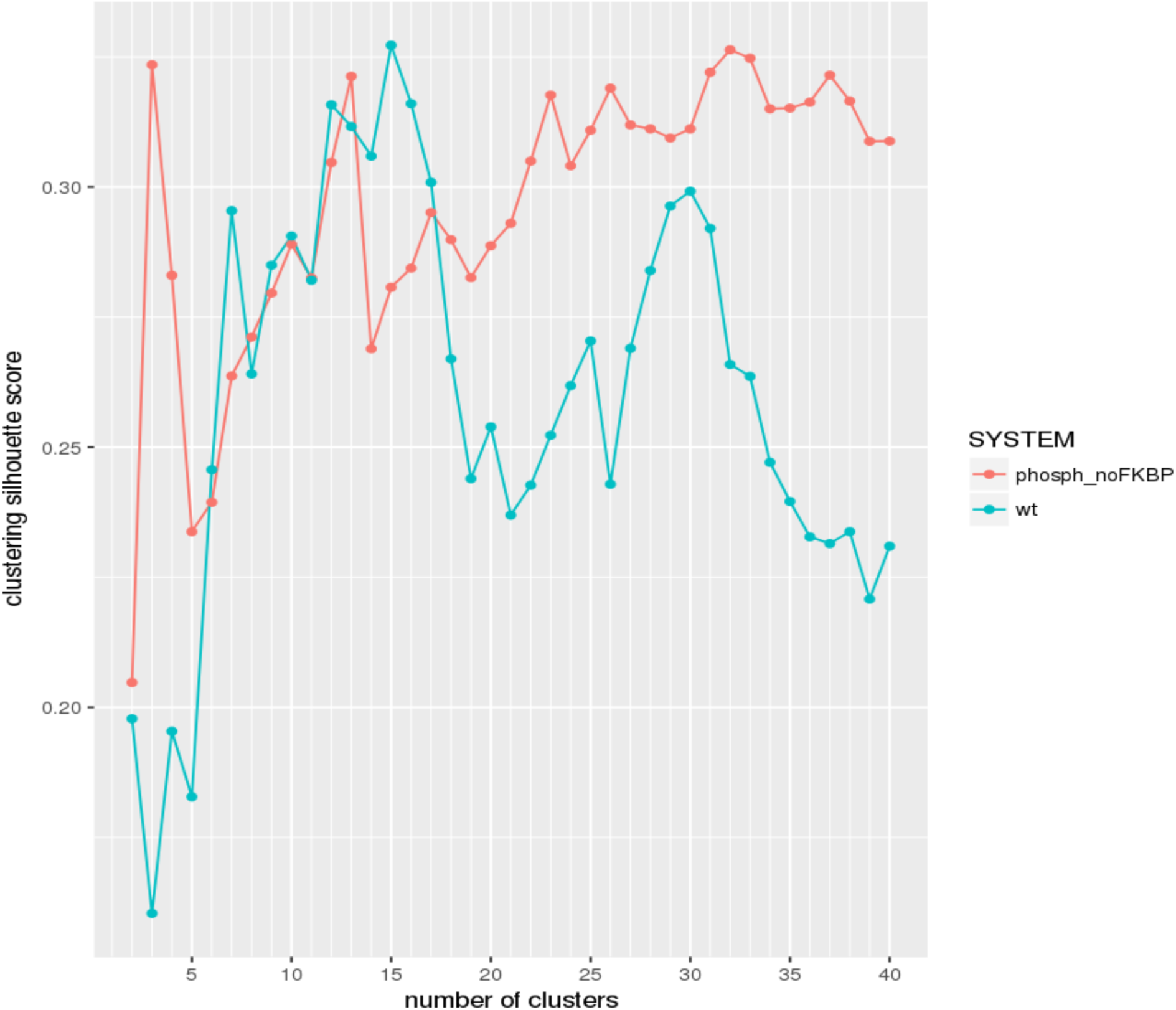
Silhouette scoring plot for temporal clustering of WAFEX data for bound unphosphorylated system (teal) and unbound phosphorylated system (red). This figure was generated by computing silhouette scores for consecutive k-cluster cuts taken from the hierarchical clustering of the frame-wise WAFEX data. Hierarchical clustering was computed using Euclidean distance norms over the log of the wavelet intensities at each frame along with Ward similarity scoring as implemented in the hclust function of the fastcluster package in R.

**Figure S6.**
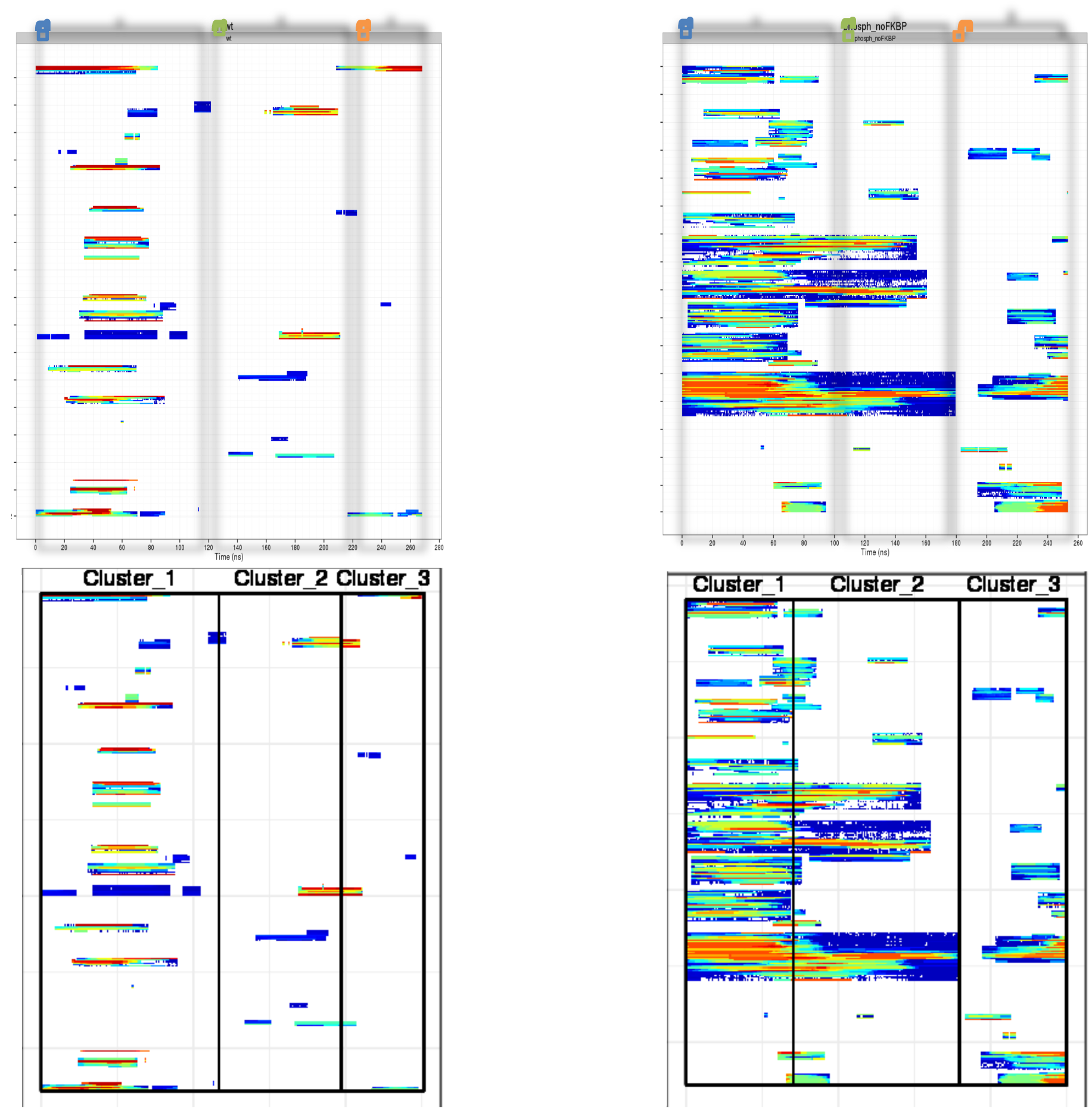
Comparison of temporal clustering of WAFEX data. Top. Clustering performed by eye. **Bottom:** Automated Hierarchical Clustering cut to 3 clusters. **Left:** Unphosphorylated ALK2 system with FKBP12 bound. **Right:** Phosphorylated ALK2 system without FKBP12 bound.

**Figure S7.**
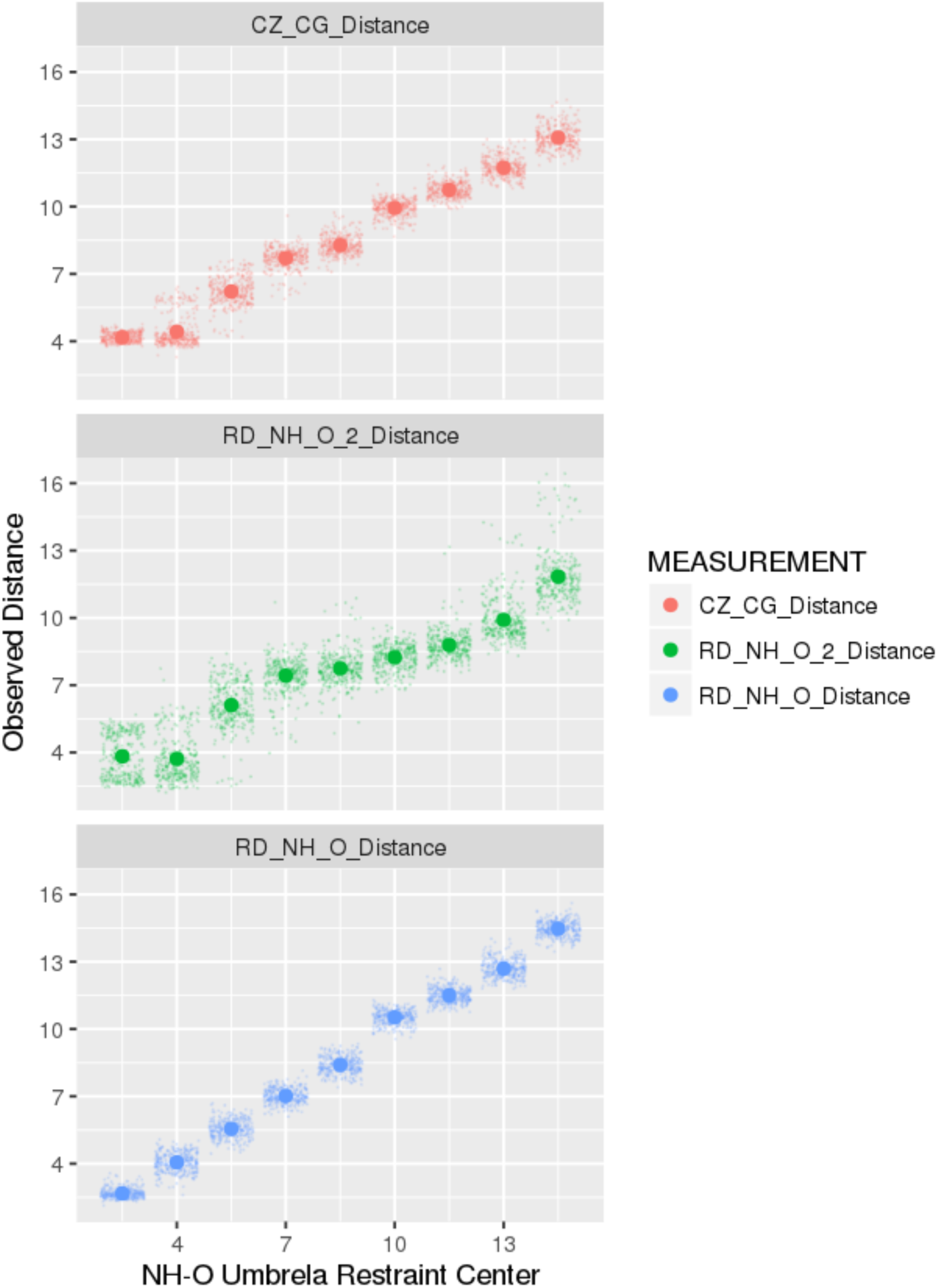
Correlation plot for R375 – D354 salt bridge distance metrics vs umbrella sampling restraint center. **Top:** Correlation plot for distance between R375(Cζ)-D354(Cγ) and umbrella restraint center. **Middle:** Correlation plot for distance between center of mass of hydrogens of non-restrained NH2 moiety of R375 guanidine group to non-restrained oxygen of D354 side-chain carboxylic acid group. **Bottom:** Correlation plot for distance between center of mass of hydrogens of restrained NH2 moiety of R375 guanidine group and restrained oxygen of D354 side-chain carboxylic acid group, i.e. umbrella restraint center.

**Movie S1. Animations of the second principal component for FKBP12-ALK2^WT^ (left, FKBP12 not shown) and ALK2^WT-Phosp^ (right) systems.** Top: Canonical kinase domain, Bot: Rotation of 90º about *z*-axis to emphasize the difference in separation between R375 and steric barrier region of αC helix.

